# A genome assembly and annotation for the Australian alpine skink *Bassiana duperreyi* using long-read technologies

**DOI:** 10.1101/2024.09.05.611471

**Authors:** Benjamin J. Hanrahan, Kirat Alreja, Andre L. M. Reis, J King Chang, Duminda S.B. Dissanayake, Richard J. Edwards, Terry Bertozzi, Jillian M. Hammond, Denis O’Meally, Ira W. Deveson, Arthur Georges, Paul Waters, Hardip R. Patel

## Abstract

The eastern three-lined skink (*Bassiana duperreyi*) inhabits the Australian high country in the southwest of the continent including Tasmania. It is an oviparous species that is distinctive because it undergoes sex reversal (from XX genotypic females to phenotypic males) at low incubation temperatures. We present a chromosome-scale genome assembly of a *Bassiana duperreyi* XY male individual, constructed using a combination of PacBio HiFi and ONT long reads scaffolded using Illumina HiC data. The genome assembly length is 1.57 Gb with a scaffold N50 of 222 Mbp, N90 of 26 Mbp, 200 gaps and 43.10% GC content. Most (95%) of the assembly is scaffolded into 6 macrochromosomes, 8 microchromosomes and the X chromosome, corresponding to the karyotype. Fragmented Y chromosome scaffolds (n=11 ≥ 1 Mbp) were identified using Y-specific contigs generated by genome subtraction. We identified two novel alpha-satellite repeats of 187 bp and 199 bp in the putative centromeres that did not form higher order repeats. The genome assembly exceeds the standard recommended by the Earth Biogenome Project; 0.02% false expansions, 99.63% kmer completeness, 94.66% complete single copy BUSCO genes and an average 98.42% of transcriptome data mappable to the genome assembly. The mitochondrial genome (17,506 bp) and the model rDNA repeat unit (15,154 bp) were assembled. The *B. duperreyi* genome assembly has one of the highest completeness levels for a skink and will provide a resource for research focused on sex determination and thermolabile sex reversal, as an oviparous foundation species for studies of the evolution of viviparity, and for other comparative genomics studies of the Scincidae.

**Species Taxonomy:** Eukaryota; Animalia; Chordata; Reptilia; Squamata; Scincidae; Lygosominae; Eugongylini; *Bassiana* (=*Acritoscincus*); *Bassiana duperreyi* (Gray, 1838) (NCBI: txid316450).

## Introduction

The family Scincidae, commonly known as skinks, is a diverse group of lizards found on all continents except Antarctica (Hedges 2014). In Australia, the Scincidae is particularly diverse, comprising 442 species in 42 genera (Cogger 2018) that occupy a wide array of habitats ranging from the inland deserts to the mesic habitats of the coast and even regions of the Australian Alps above the snowline. The eastern three-lined skink (*Bassiana duperreyi* (Gray 1838), sensu Hutchinson *et al*. 1990) is a species complex in the Eugongylus group of Australian Lygosominae skinks that is found in the south of eastern Australia, including Tasmania and islands of Bass Strait. The alpine taxon within this species complex, as defined by mitochondrial (Dubey and Shine 2010) and nuclear DNA sequence variation (Dissanayake *et al*. 2022), occupies the highlands and alpine regions of the states of New South Wales, Victoria and Tasmania. It is hereafter referred to as the Alpine three-lined skink (Figure 1). The alpine taxon is genetically distinct from other members of the species complex that occupy the lowlands and coastal regions of Victoria and South Australia, the two of which probably represent distinct species (Dissanayake *et al*. 2022). We report on the genome assembly and annotation for the Alpine clade of the three-lined skink (Figure 1c).

**Figure 1.**
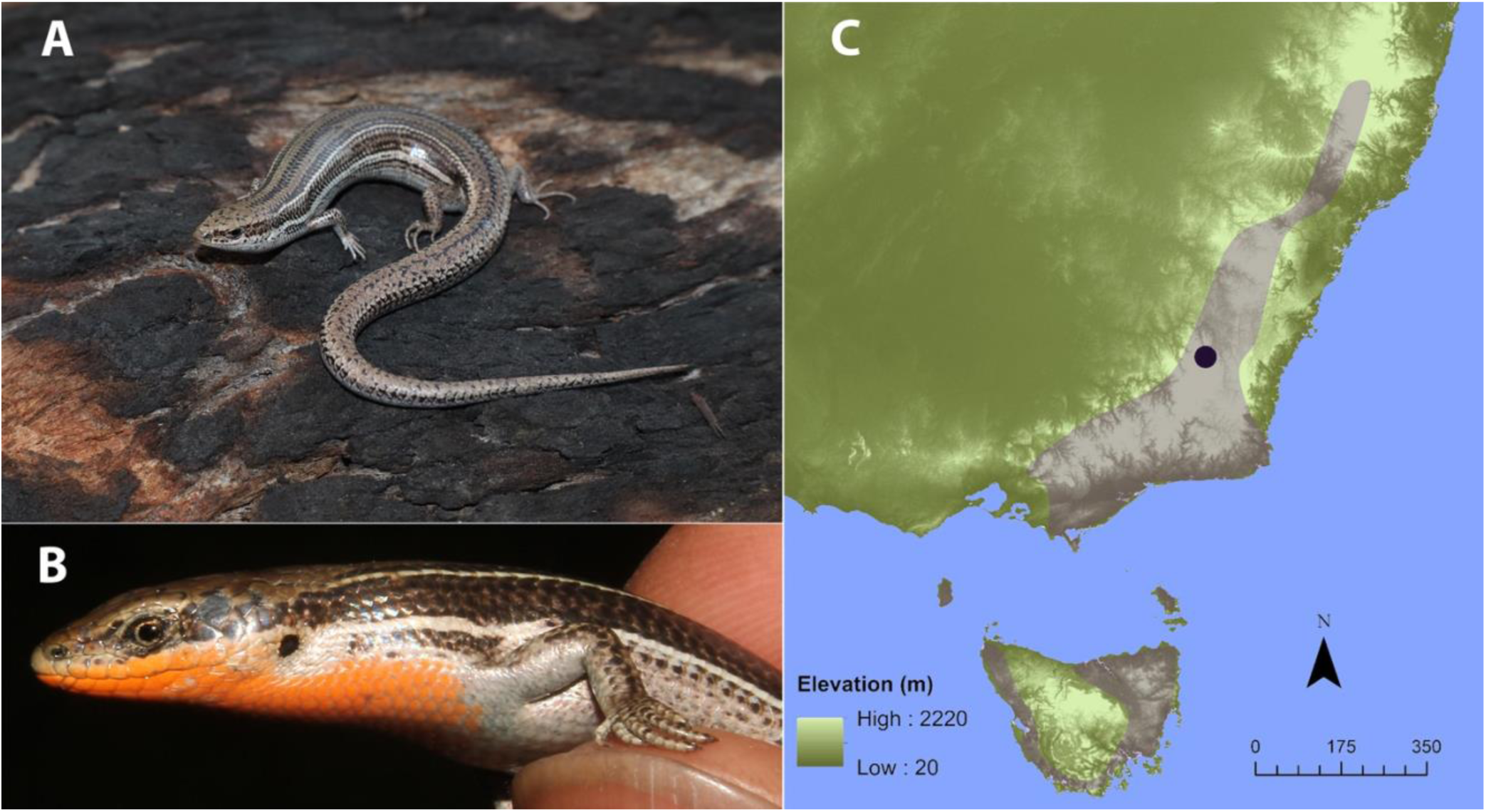
The Alpine three-lined skink *Bassiana duperreyi* from the Brindabella Range, Australian Capital Territory. A: Representative female of the species; B: Male dividual (DDBD_364) that was sequenced for the genome assembly and annotation, showing the distinctive ventral breeding colour; C: Distribution of the Alpine three-lined skink shown in gray (after Dissanayake *et al*. 2022). Location of collection of the focal male shown as a black dot.

*Bassiana duperreyi* has well-differentiated sex chromosomes and male heterogamety (XX/XY) with 6 macrochromosome pairs, 8 microchromosome pairs and a sex chromosome pair (2n=30, Figure 3, Dissanayake *et al*. 2020). The taxon is interesting from a genomic perspective because there are relatively few genome assemblies for this very diverse group of lizards, and because candidates for the sex determination gene in reptiles with genetic sex determination are few and poorly characterized (Deakin *et al*. 2016; Zhang *et al*. 2022). Additionally, the developmental program initiated by genetic sex determination can be diverted by low temperature incubation in the laboratory and in the wild (Radder *et al*. 2008; Holleley et al 2016; Dissanayake *et al*. 2021a,b; Dissanayake 2022). Sex determination and sex reversal is a major focus for research on this species. *Bassiana duperreyi* is also of interest because it is oviparous, serving as a foundational model for understanding viviparity and placentation in other species within the subfamily Eugongylinae of Lygosomatine skinks (Stewart and Thompson 1996), and it is recognized as a significant contributor to the study of reproductive biology among Australian lizards (Van Dyke *et al*. 2021).

**Figure 2.**
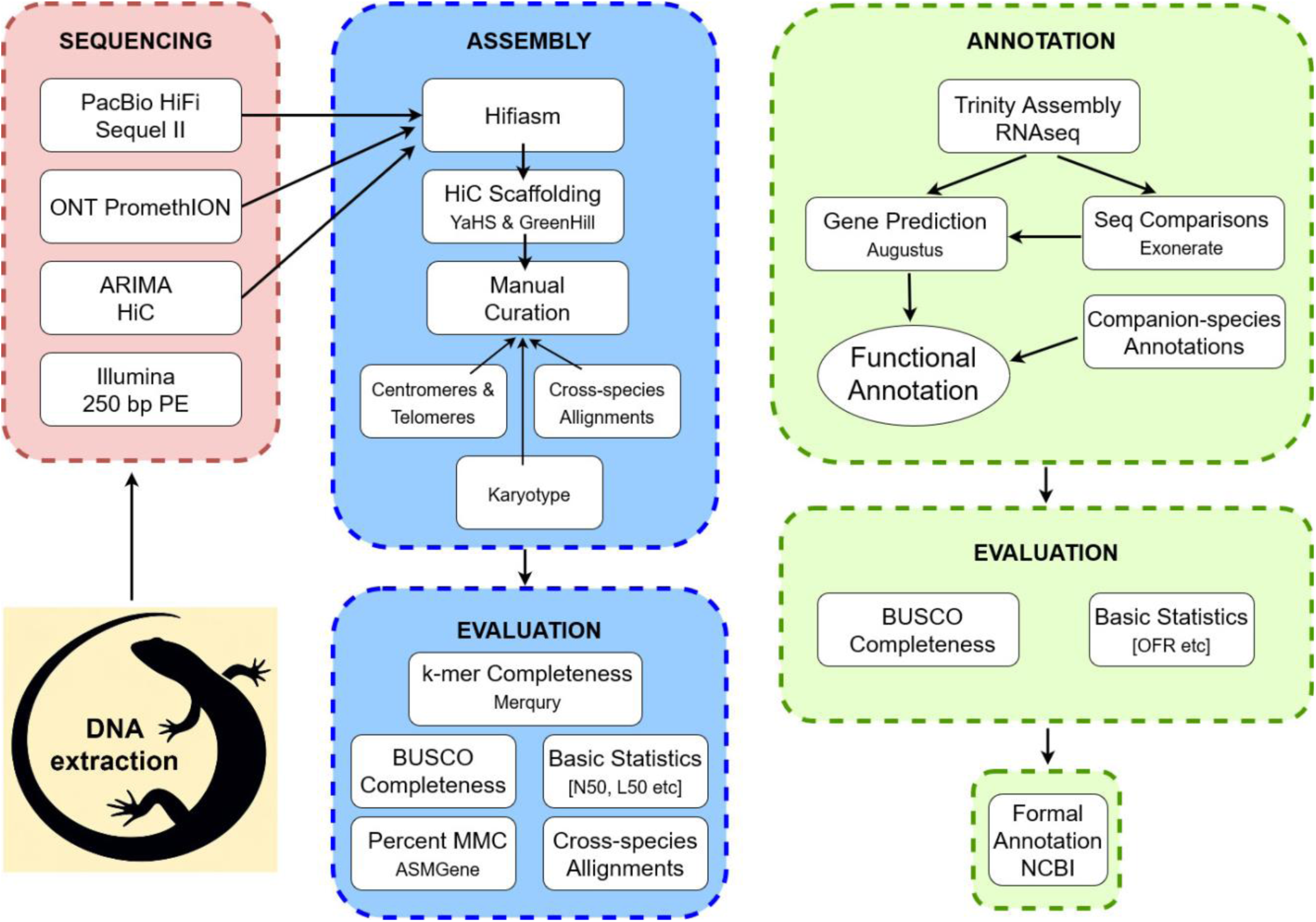
Schematic overview of the JigSaw workflow for sequencing, assembly and annotation of the *B. duperreyi* genome. Illumina 250 bp PE reads were initially generated to polish the ONT reads, no longer necessary because of increases in the accuracy of ONT reads, and for the identification of Y-enriched kmers. They have been used for quality assessment of the genome and genome subtraction. Steps employed for quality control of sequence data not shown. Repeat annotation was undertaken with Repeatmasker.

**Figure 3.**
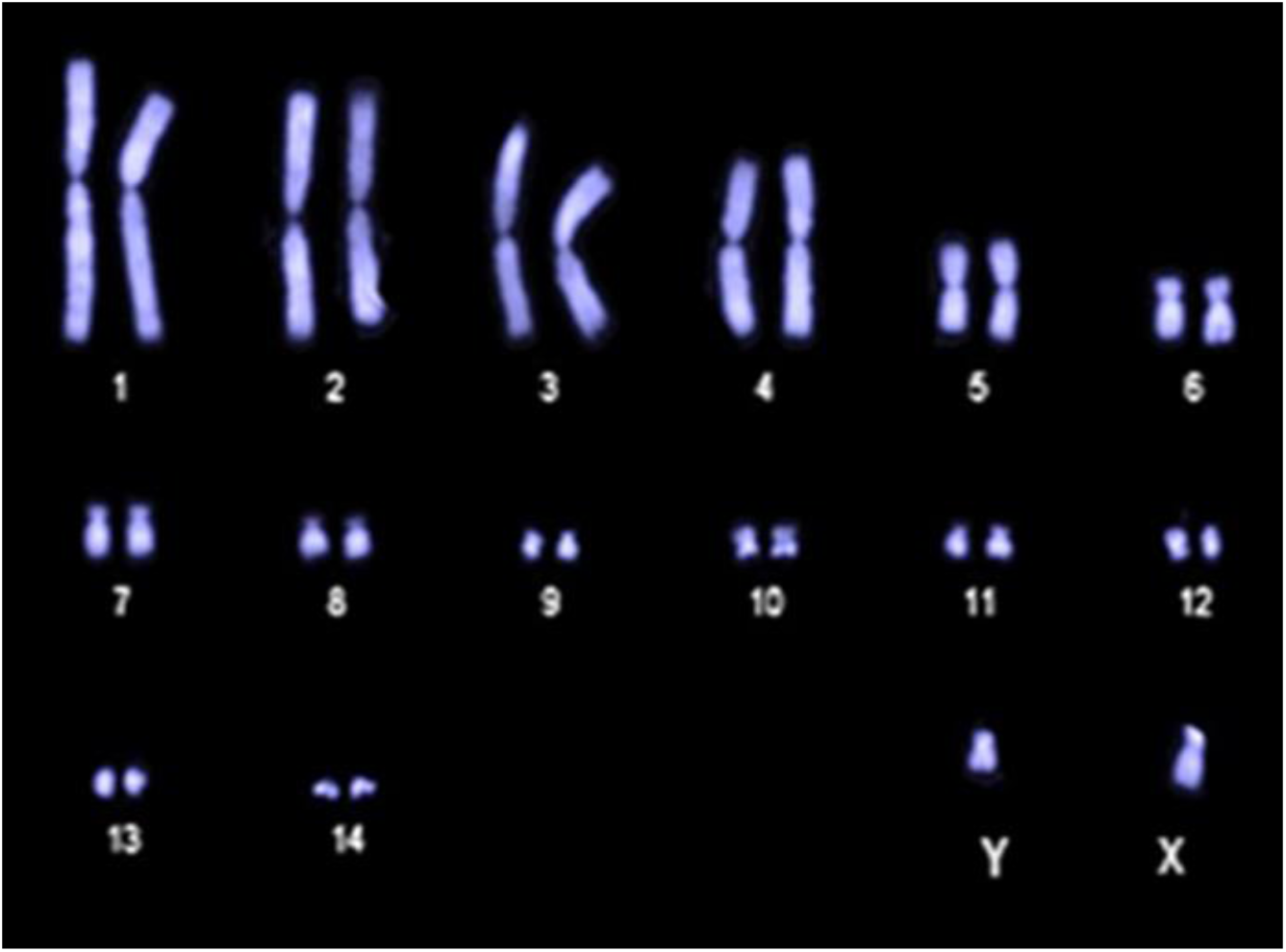
Karyotype for *Bassiana duperreyi* (SpecimenDDBD_142 XY male, Piccadilly Circus, Brindabella Range, ACT –35.361658 148.803458) [after Dissanayake *et al*. 2020]. Chromosome number: 2n=30.

Research in these areas of interest will be greatly facilitated by a high-quality draft genome assembly for *B. duperreyi*. The ability to generate telomere to telomere assemblies and identify the non-recombining regions of the sex chromosomes, within which lies any master sex determining gene, will greatly narrow the field of candidate sex determining genes in skinks. Furthermore, the disaggregation of the X and Y (or Z and W) sex chromosome haplotypes (phasing) will allow comparison of the X and Y sequences to gauge putative loss or gain of function in key sex gene candidates. In studies of the evolution of viviparity of model species such as the Australian tussock cool-skink *Pseudemoia entrecasteauxii* (Adams *et al*. 2005), a high-quality genome assembly for a closely-related oviparous species such as *B. duperreyi* provides a basis for comparisons of transcriptional profiles of putative genes governing reproduction and related studies of differential gene family proliferation (Griffith *et al*. 2016).

In this paper, we present an annotated assembly of the genome of the Alpine three-lined skink *Bassiana duperreyi* as a resource to enable and accelerate research into the unusual reproductive attributes of this species and for comparative studies across the Scincidae and reptiles more generally.

## Materials and Methods

Software and databases used in this paper are provided with version numbers, URL links and citations in Table S1.

### Sample collection

The focal male individual for the *B. duperreyi* genome assembly was collected from Mt Ginini in the Brindabella Ranges, Australia (–35.525S 148.783E, Figure 1c). A detailed description of the study site is available (Dissanayake *et al*. 2022). Phenotypic sex was determined by hemipene eversion (Harlow 1996) and by conspicuous male breeding coloration (Figure 1b). The individual was transported to the University of Canberra and euthanised. Tissue and blood samples were collected and snap frozen in liquid nitrogen. An additional blood sample was preserved on a Whatman FTA™ Elute Card (WHAWB12-0401, GE Healthcare UK Limited, UK). DNA was extracted from the FTA™ Elute Card for a sex test based on PCR to confirm chromosomal sex as XY (Dissanayake *et al*. 2020).

### DNA Extraction and Sequencing

Sequencing data were generated using four platforms: Illumina® short-read platform, PacBio HiFi and Oxford Nanopore Technologies (ONT) long-read platforms and HiC linked-reads using the Arima Genomics platform (Figure 2). *Illumina sequence data:* Genomic DNA was extracted from muscle tissue using the salting out procedure (Miller *et al*. 1988). Sequencing libraries were prepared using Illumina DNA PCR-Free Prep library kit and sequenced on the Illumina NovaSeq instrument in 250 bp paired-end format with *ca* 500 bp fragment size. DNA quality assessments, library preparation and sequencing were performed by the Ramaciotti Centre for Genomics (UNSW, Sydney, Australia). Summary statistics for the Illumina data are provided in Table S2.

*PacBio HiFi sequence data:* Genomic DNA was extracted from muscle tissue using the salting out procedure (Miller et al. 1988) and spooled to enrich for high molecular weight DNA. Sequencing libraries were prepared and sequenced on PacBio Sequel II machine using two SMRTCells as per the manufacturer’s protocol. The Australian Genomics Research Facility (AGRF), Brisbane, Australia, performed DNA quality assessment, library preparation and sequencing. *DeepConsensus* (v1.2.0, Baid *et al*. 2023) was used to perform base calling from subreads. Subsequently, *Cutadapt* (v3.7, Martin *et al*. 2011, parameters: error-rate 0.1 –overlap 25 –match-read-wildcards –revcomp –discard-trimmed) was used to remove reads containing PacBio adapter sequences to obtain analysis-ready sequence data. Quality statistics are provided in Figures S1 and additional statistics in Table S3.

*ONT sequence data:* Genomic DNA was extracted from 13 mg of ethanol-preserved muscle tissue, using the Circulomics Nanobind tissue kit (PacBio, Menlo Park, California) as per the manufacturer’s protocols, including the specified pre-treatment for ethanol removal. Library preparation was performed with 3 µg of DNA as input, using the SQK-LSK109 kit from Oxford Nanopore Technologies (Oxford, UK) and sequenced across two promethION (FLO-PRO002, R9.4.1) flow cells, with washes (EXP-WSH004) performed every 24 hr. ONT signal data was converted to *slow5* format using *slow5tools* (v1.1.0, Samarakoon *et al*. 2023b) and base calling was performed using Oxford Nanopore’s basecaller *dorado* (v7.2.13) and *buttery-eel* (v0.4.2, Samarakoon *et al*. 2023a) wrapper scripts. Parameters were chosen to remove adapter sequence *(--detect_mid_strand_adapter –– trim_adapters ––detect_adapter ––do_read_splitting)* and the super accuracy “*dna_r9.4.1_450bps_sup.cfg*” model was used for base calls. Quality statistics are provided in Figures S1, and additional statistics in Table S4.

*Arima Genomics HiC sequence data:* A liver sample was processed for HiC library preparation and sequencing by the Biological Research Facility (BRF) at the Australian National University using the Arima Genomics HiC 2.0 kit (Carlsbad, California). The library was sequenced across two lanes of the Illumina S1 flowcell on NovaSeq 6000 machine in 150 bp paired-end format. Summary statistics are provided in Table S5.

*Transcriptome sequence data:* We used transcriptome sequence from a larger cohort of 30 male and female animals to develop gene models for the assembly. Total RNA was extracted from the brain, heart, ovary, testis (“DDBD” prefix, Table S6) by the Garvan Molecular Genetics unit (Sydney). We included other sequences previously generated in our laboratory but unpublished (“DOM” prefix, Table S6 from brain, liver, testes, ovary) and sequences from10 uterine samples (“BD” prefix, Table S6, Foster *et al*. 2022). Briefly, tissue extracts were homogenized using T10 Basic ULTRA-TURRAX® Homogenizer (IKA, Staufen im Breisgau, Germany), RNA was extracted using TRIzol reagent (Thermo Scientific, Waltham, Massachusetts) following the manufacturer’s instructions, and purified by isopropanol precipitation. Seventy-five bp single-end reads were generated for recent samples on the Illumina NextSeq 500 platform at the Ramaciotti Centre for Genomics (UNSW, Sydney, Australia). Some earlier libraries were sequenced with 100 bp PE reads.

### Karyotype

The karyotype for the alpine form of *B. duperreyi* was obtained from the supplementary material accompanying Dissanayake *et al*. (2020) (Figure 3) to provide an expectation for final telomere to telomere scaffolding by the assembly.

### Assembly

All data analyses were performed on the high-performance computing facility, Gadi, hosted by Australia’s National Computational Infrastructure (NCI, https://nci.org.au). Scripts are available at https://github.com/kango2/ausarg.

#### Primary genome assembly

PacBio HiFi, ONT and HiC sequence data were used to generate interim haplotype consensus and haplotype assemblies using *hifiasm* (v0.19.8, Cheng et al. 2021, 2022, default parameters). HiC data were aligned to the interim haplotype consensus assembly using the *Arima Genomics alignment pipeline* following the user guide. HiC read alignments were processed using *YaHS* (v1.1, Zhou et al. 2022, parameters: –r 10000, 20000, 50000, 100000, 200000, 500000, 1000000, 1500000) to generate scaffolds. Range resolution parameter (-r) in *YaHS* was restricted to 1500000 to ensure separation of microchromosomes into individual scaffolds. Vector contamination was assessed using *VecScreen* defined parameters for *BLAST* (v2.14.1, parameters: –task blastn –reward 1 –penalty –5 –gapopen 3 –gapextend 3 –dust yes –soft_masking true –evalue 700 –searchsp 1750000000000) and the *UniVec* database (accessed on 18^th^ June 2024). Putative false expansion and collapse metrics were calculated using the *Inspector* (v1.2, default parameters) and PacBio HiFi data.

#### Read depth and GC content calculations

PacBio HiFi (parameter: –x map-pb) and ONT (parameter: –x map-ont) sequence data were aligned to the scaffold assembly using *minimap2* (v2.17, Li 2018) Similarly, Illumina sequence data were aligned to the assembly using *bwa-mem2* (v2.2.1, Vasimuddin *et al*. 2019) using default parameters. Resulting alignment files were sorted and indexed for efficient access using *samtools* (v1.19, Danecek *et al*. 2021). Read depth in non-overlapping sliding windows of 10 Kbp was calculated using the *samtools bedcov* command. GC content in non-overlapping sliding windows of 10 Kbp was calculated using *calculateGC.py* script.

#### Centromeric alpha satellite and telomere repeats

*TRASH* (v1.12, Wlodzimierz et al., 2023, parameters: –N.max.div 5) was used to identify putative satellite repeat units. Repeat units spanning >100 Kbp were prioritized to detect putative centromeric satellite repeat motifs. Two unique repeat motifs with monomer period sizes of 199 bp and 187 bp were identified and labeled as centromeric satellite repeats. These two motifs were supplied to the *TRASH* as templates for refining the centromeric satellite repeat annotations. For telomeric repeat detection, *Tandem Repeat Finder* (*TRF*) (v4.09.1, Benson 1999, parameters: 2 7 7 80 10 500 6 –l 10 –d –h) was used to detect all repeats up to 6 bp length. TRF output was processed using *processtrftelo.py* script to identify regions >600 bp that contained conserved vertebrate telomeric repeat motif (TTAGGG). These regions were labeled as potential telomeres.

#### Sex chromosome assembly

Scaffolds associated with the sex chromosomes were identified using read depth. The putative X scaffold will have half the read depth of the autosomal scaffolds in an XY individual. The Y chromosome scaffolds were identified by a process of elimination, removing scaffolds already assigned to large scaffolds with read depths corresponding to the genome average, and removing scaffolds that were associated primarily with rDNA or centromeric satellite repeats. Y enriched contigs, obtained by genome subtraction (Dissanayake *et al*. 2020), were mapped to the remaining scaffolds and those with a high density of mapped contigs were considered to be Y chromosome scaffolds.

#### Mitochondria genome assembly

PacBio HiFi sequence data were used to assemble and annotate mitochondrial genome using *mitoHiFi (v3.2.2,* Uliano-Silva *et al*. 2023). Mitochondrial genome (NCBI Accession: NC_066473.1, Wu *et al*. 2022) of the Hainan water skink, *Tropidophorus hainanus*, was used as a reference for *mitoHiFi*. The mitochondrial genome of *B. duperreyi* was aligned to scaffolds using *minimap2 (-x asm20)* to identify and remove erroneous mitochondrial scaffolds and retain a single mitochondrial genome sequence.

#### Manual editing of scaffolds

Read depth, GC content, and centromere and telomere locations for *YaHS* scaffolds >1 Mbp length were visually inspected. Three scaffolds contained internal telomeric repeat sequences near the *YaHS* joined contigs (Figure S2), which were interpreted as false-positive joins by *YaHS* scaffolder and were subsequently split at the gaps using *agptools*.

### Assembly evaluation

#### RNAseq mapping rate

RNAseq data from multiple tissues (Table S6) were aligned to the assembly using *subread-align* (v2.0.6, Liao *et al*. 2013) to calculate percentage of mapped fragments for evaluating RNAseq mapping rate.

#### Assembly completeness and per base error rate estimation

Illumina sequence data were trimmed for adapters and low-quality using *Trimmomatic* (v0.39, Bolger *et al*. 2014, parameters: ILLUMINACLIP:TruSeq3-PE.fa”:2:30:10:2:True LEADING:3 TRAILING:3 SLIDINGWINDOW:4:20 MINLEN:36). Resultant paired-end sequences were used to generate kmer database using *meryl* (v1.4.1, Rhie *et al*. 2020). Merqury (v1.3, Rhie et al. 2020) was used with *meryl* kmer database to evaluate assembly completeness and estimate per base error rate of pseudo-haplotype and individual haplotype assemblies.

#### Gene completeness evaluation

*BUSCO* (v5.4.7, Manni *et al*. 2021) was run using *sauropsida_odb10* library in offline mode to assess completeness metrics for conserved genes. BUSCO synteny plots were created with *ChromSyn* (v1.3.0, Edwards et al. 2022).

### Annotation

#### Repeat annotation

*RepeatModeler* (v2.0.4, parameters: –engine ncbi) was used to identify and classify repetitive DNA elements in the genome. Subsequently, *RepeatMasker* (v4.1.2-pl) was used to annotate and soft-mask the genome assembly using the species-specific repeats library generated by *RepeatModeler* and families were labelled accordingly.

#### Ribosomal DNA

Assembled scaffolds were aligned to the 18S small subunit (n=1,415) and 28S large subunit (n=283) sequences of deuterostomes obtained from the SILVA ribosomal RNA database (v138.1, Quast *et al*. 2013) using minimap2 (v2.26, Li 2018, parameters: ––secondary=no). Alignments with >50% bases covered for 18S and 28S subunits were retained. These scaffolds were labelled as rDNA scaffolds.

#### De novo gene annotations

RNAseq data from multiple tissues (Table S6) were processed using *Trinity* (v2.12.0, Grabherr *et al*. 2011, parameters: ––min_kmer_cov 3 ––trimmomatic) to produce individual transcriptome assemblies. Parameters were chosen to remove low abundance and sequencing error k-mers. The assembled transcripts were aligned to the UniProt-SwissProt database (last accessed on 28-Feb-2024) using *diamond* (v2.1.9, Buchfink *et al*. 2021, parameters: blastx ––max-target-seqs 1 –iterate ––min-orf 30). Alignments were processed using *blastxtranslation*.*pl* script to obtain putative open reading frames and corresponding amino acid sequences. Transcripts containing both the start and the stop codons, with translated sequence length between 95% and 105% of the best hit to UniProt_SwissProt sequence, were selected as full-length transcripts.

Amino acid sequences of full-length transcripts were processed using *CD-HIT* (v4.8.1, Fu *et al*. 2012, parameters: –c 0.8 –aS 0.9 –g 1 –d 0 –n 3) to cluster similar sequences with 80% pairwise identity and where the shorter sequence of the pair aligned at least 90% of its length to the larger sequence. A representative transcript from each cluster was aligned to the repeat-masked genome using *minimap2* (v2.26, parameters: ––splice:hq), and alignments were coordinate-sorted using *samtools*. Transcript alignments were converted to *gff3* format using *AGAT* (v1.4.0, agat_convert_minimap2_bam2gff.pl) and parsed with *genometools* (v1.6.2, Gremme *et al*. 2013) to generate training gene models and hints for *Augustus* (v3.4.0, Stanke et al. 2008) with untranslated regions (UTRs). Similarly, transcripts containing both start and stop codons with translated sequence length outside of 95% and 105% of the best hit to UniProt_SwissProt sequence, were processed in the same way to generate additional hints. A total of 500 of these representative full-length transcripts were used in training for gene prediction to calculate species-specific parameters. During the gene prediction model training, parameters were optimized using all 500 training gene models with a subset of 200 used only for intermediate evaluations to improve run time efficiency. Gene prediction for the full dataset used 20 Mbp chunks with 2 Mbp overlaps to improve run time efficiency. Predicted genes were aligned against Uniprot_Swissprot database for functional annotation using best-hit approach and *diamond*. Unaligned genes were subsequently aligned against Uniprot_TrEMBL database for functional annotation. The quality of the final assembly was assessed using various standard measures (Figure 2) as described by the Earth Biogenomes Project (EBP, https://www.earthbiogenome.org/report-on-assembly-standards, Version 5).

### Other

Common names for species referred to are as follows: Australian blue-tongued lizard *Tiliqua scincoides,* African cape cliff lizard *Hemicordylus capensis,* Australian olive python *Liasis olivaceus,* cobra *Naja naja,* Prairie rattlesnake *Crotalus viridis,* Chinese crocodile lizard *Shinisaurus crocodilurus,* green anole *Anolis carolinensis,* Madagascan panther chameleon *Furcifer pardalis,* European sand lizard *Lacerta agilis,* Binoe’s gecko *Heteronotia binoei,* and leopard gecko *Eublepharis macularius*.

## Results and Discussion

### DNA sequence data quantity and quality

PacBio HiFi sequencing yielded 52.4 Gb with a median read length of 14,962 bp (Table 1) and 82.1% of reads with mean quality value Q30. Similarly, ONT sequencing yielded 104.5 Gb with an N50 value of 10,945 bp and 50.4% reads with mean quality value Q20. Illumina sequencing in 250 bp paired-end format yielded 110.6 Gb sequence data and HiC yielded 81.8 Gb sequence data. The distributions of quality scores and read lengths for the long-read sequencing align with known characteristics of the ONT and PacBio platforms (Figure S1). K-mer frequency histograms of Illumina, ONT and PacBio HiFi sequence data for k=17, k=21 and k=25 show two distinct peaks (Figure 4) confirming the diploid status of this species. The peak for heterozygous k-mers was smaller for k=17 compared to the homozygous k-mer peak. In contrast, the heterozygous k-mer peak was higher for k=25 compared to the homozygous k-mer peak, suggestive of high heterozygosity at a small genomic distance. Genome size was estimated to be 1.64 Gb using the formulae of Georges *et al*. (2015) and Illumina sequence data, with a k-mer length of 17 bp, homozygous peak of 63 (Figure 4) and the mean read length of 241.2 bp. Read depth, obtained by dividing the total DNA sequence data from each platform by the genome size, was consistent with that typically generated by PacBio HiFi and Illumina platforms respectively (Table 1). Assembly sizes were consistent with the estimates of median read depths of 64.84*x* for ONT, 34.49*x* PacBio HiFi and 71.40*x* Illumina platforms calculated for 10 Kbp non-overlapping sliding windows of the assembly.

**Figure 4.**
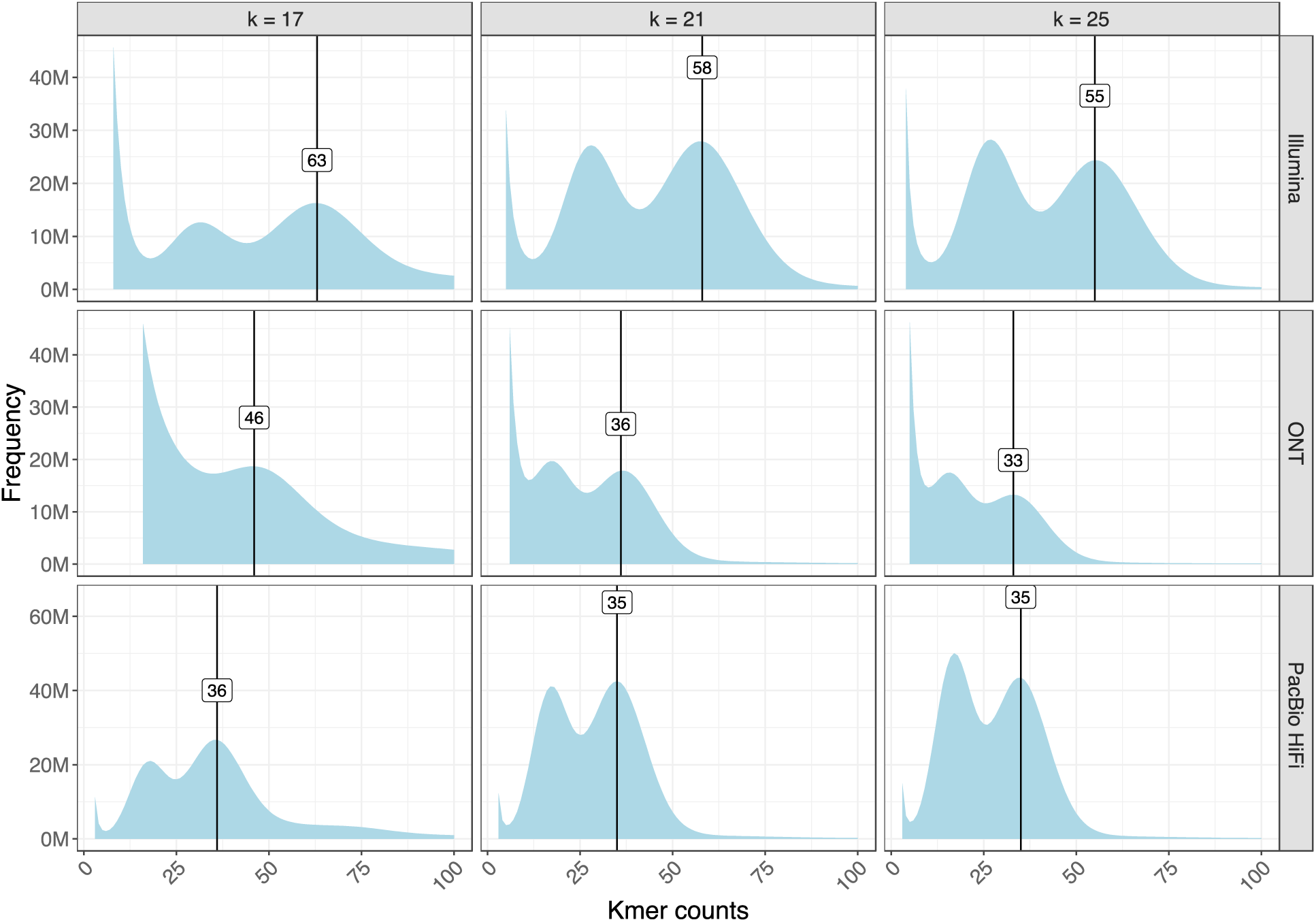
Distribution of k-mer counts using sequences from Illumina, Oxford Nanopore Technologies (ONT), and PacBio (PB) platforms for *Bassiana duperreyi*. Heterozygosity is high as indicated by dual peaks in each graph, and the height of the heterozygous peak increases with the length of the k-mer. This confirms diploidy.

**Table 1.** Summary metrics for sequence data and assembly for *Bassiana duperreyi*.

| Sequencing Platform | Number of Reads | Mean Read Length (bp) | Median Read Length (bp) | Total Bases | Estimated Read Depth |
| --- | --- | --- | --- | --- | --- |
| Illumina PE DNA | 458,637,888 | 241.2 | 241 | 110,612,868,725 | 70.55x |
| PacBio HiFi Sequel II | 3,395,376 | 15,443 | 14,962 | 52,437,383,684 | 33.44x |
| ONT R9.4.1 | 22,044,338 | 4,739 | 2,114<br>(n50 = 10,945) | 104,472,064,570 | 66.63x |
| Arima Genomics HiC | 270,940,642 | 151 | 151 | 81,824,073,884 | -- |

### Assembly

Hifiasm produced three assemblies: one for each haplotype and a haplotype consensus assembly of high quality as evidenced by assembly metrics (Table 2). The haplotype consensus assembly was chosen for further scaffolding using the HiC data to improve assembly contiguity, and then manually curated (Figures S2, S6). Scaffold numbers 7, 10 and 13 were split at internal telomere sequences (Figure S2). Scaffolding markedly improved contiguity of the assembly presented here. The final reference genome for *B. duperreyi* had a total length of 1,567,894,183 bp assembled into 172 scaffolds, with 54 gaps each marked by 200 Ns, which compares well with other published squamate genome assemblies (Table S7). The assembly size of 1.57 Gb is 71.4 Mb shorter than the expected genome size. This is likely because of the collapse of ribosomal DNA copies, satellite repeat units of centromeres and the Y chromosome, and heterozygous indels. There were 68 regions of >50 bp length spanning 41,549 bp identified as putatively collapsed and 240 regions spanning 309,329 bp (0.02% of the assembly length) as putative expansions.

**Table 2.**
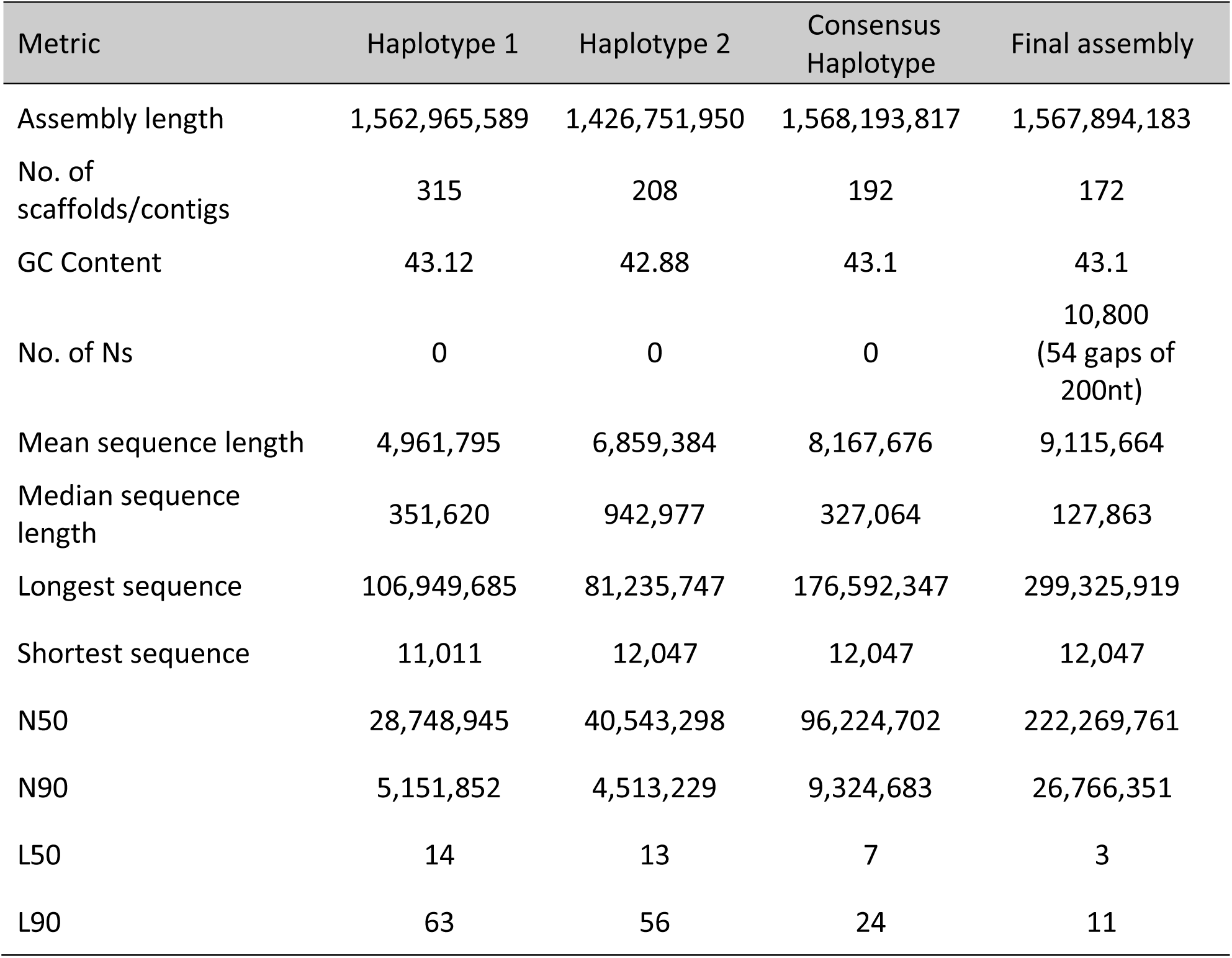
Summary metrics for the genome assembly of *Bassiana duperreyi*. Refer to Table S7 for comparisons with other species.

| Metric | Haplotype 1 | Haplotype 2 | Consensus Haplotype | Final assembly |
| --- | --- | --- | --- | --- |
| Assembly length | 1,562,965,589 | 1,426,751,950 | 1,568,193,817 | 1,567,894,183 |
| No. of scaffolds/contigs | 315 | 208 | 192 | 172 |
| GC Content | 43.12 | 42.88 | 43.1 | 43.1 |
| No. of Ns | 0 | 0 | 0 | 10,800<br>(54 gaps of 200nt) |
| Mean sequence length | 4,961,795 | 6,859,384 | 8,167,676 | 9,115,664 |
| Median sequence length | 351,620 | 942,977 | 327,064 | 127,863 |
| Longest sequence | 106,949,685 | 81,235,747 | 176,592,347 | 299,325,919 |
| Shortest sequence | 11,011 | 12,047 | 12,047 | 12,047 |
| N50 | 28,748,945 | 40,543,298 | 96,224,702 | 222,269,761 |
| N90 | 5,151,852 | 4,513,229 | 9,324,683 | 26,766,351 |
| L50 | 14 | 13 | 7 | 3 |
| L90 | 63 | 56 | 24 | 11 |

The *Bassiana duperreyi* genome is contiguous with a scaffold N50 value of 222,269,761 bp and a N90 value of 26,766,351 with the largest scaffold of 299,325,919 bp (Table 2). L50 and L90 values were 3 and 11 respectively, typical of species with microchromosomes, where most of the genome is present in large macrochromosomes.

Of the 15 major scaffolds in the *YaHS* assembly (corresponding in number to the chromosomes in the karyotype of *B. duperreyi*, Figure 3), each had a single well-defined centromere. Seven were complete in the sense of having a single centromere and two terminal telomeric regions (Figure 5). A further 6 were missing one telomeric region and 2 were missing telomeres altogether. Telomeres were comprised of the vertebrate telomeric motif TTAGGG and ranged in size from the minimum threshold of 100 copies to ca 3,200 copies (BASDUscf12). The telomeric regions were typically characterized by an expected rise in GC content (Figure 5). Centromeric repeats comprised two repeat families, one based on a motif 199 bp in length (CEN199) and restricted to the centromeric region. The other was based on a motif 187 bp in length (CEN187) that was found both within and outside the centromeric region (Figure 5). Refer to Table S8 for the sequences and their coordinates and Table S9 for repeat counts. The centromeric repeat regions were characterized by a drop in read depth, arising from difficulties in mapping reads in those regions, and by a drop in GC content that was most pronounced in the CEN199 repeats (Figure 5).

**Figure 5.**
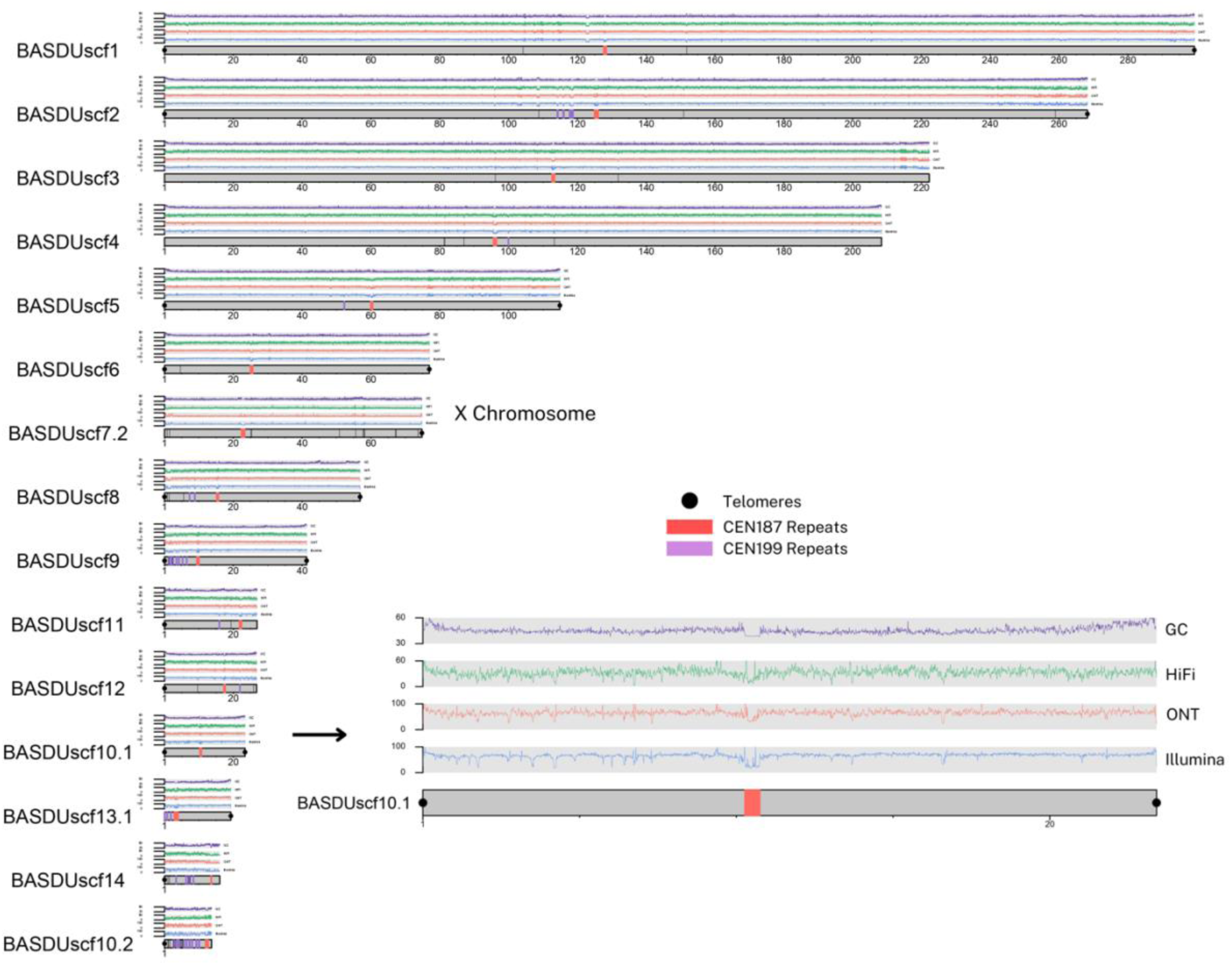
A plot of the 15 longest scaffolds (corresponding to the number of chromosomes of *Bassiana duperreyi*) for the*YaHS* assembly. The Y chromosome was fragmented (n = 21 fragments, 11 ≥ 1 Mbp) and not shown (refer Figure S3). Four traces are shown. The top trace (purple, range 30-60%) represents GC content, the next trace (Green, range 0-50*x*) represents PacBio HiFi read depth, the next trace (red, range 0-100*x*) represents Oxford Nanopore PromethION read depth, and the fourth trace (blue, range 0-100*x*) represents Illumina read depth. The inset shows Scaffold BASDUscf10.1 is enlarged for illustration. Note that centromeric sequence (red bars, CEN199; purple bars, CEN187) was often associated with a distinct drop in GC content and read depth. Black dots indicate telomeric sequence. Refer to the https://github.com/kango2/basdu for a high-resolution version of this figure.

### Assembly evaluation

Completeness of the assembly was estimated to be 88.32% and the per base assembly quality estimate was 56.54 (1 error in 221,986 bp). High heterozygosity in the k-mer profiles affects assembly completeness metrics measured by *Merqury*. Individual haplotype assemblies were 88.21% and 84.38% complete, which as expected was similar to that of the consensus haplotype assembly. However, of all the assessable k-mers by *Merqury*, 99.63% were present in one of the two haplotypes (Figure 6). This shows that assembly completeness metrics for a consensus haplotype assembly measured using k-mers can be understated for species with high heterozygosity.

**Figure 6.**
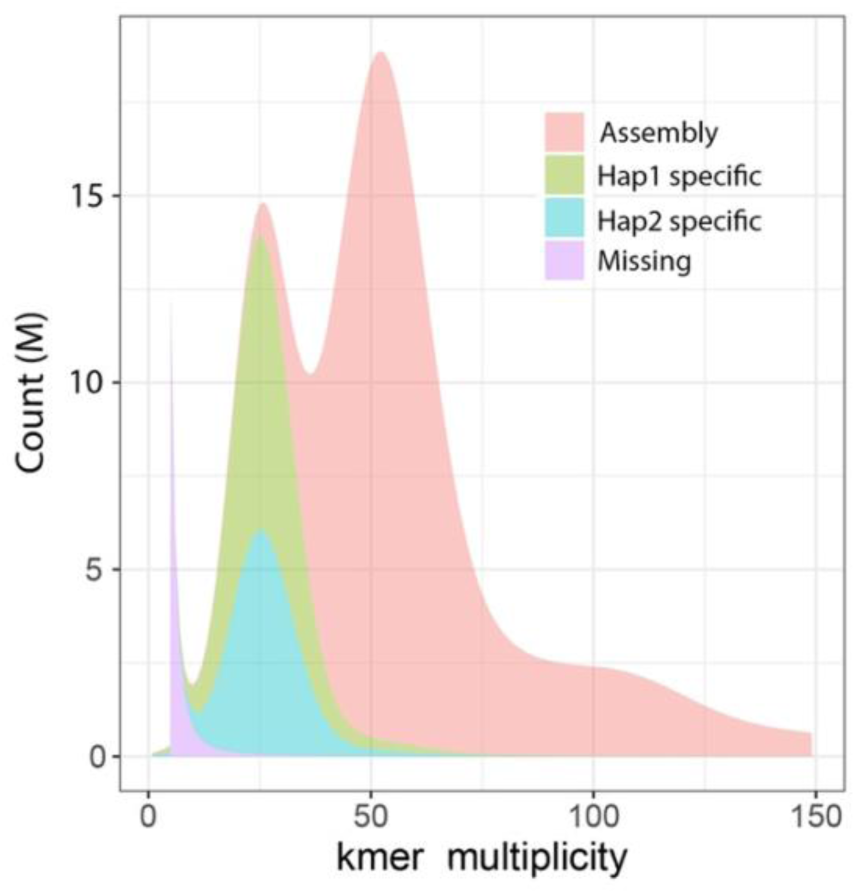
Distribution of Illumina k-mers (k = 17) in the genome assembly of *Bassiana duperreyi*. K-mer counts are shown on the x-axis and the frequency of occurrence of those counts on the y-axis. Those scored as missing are found in reads only.

Analyses using the Benchmarking Universal Single-Copy Orthologs (BUSCO) gene set for Sauropsids reveals 94.70% genes as complete, with a minimal proportion duplicated (D:2.4%), indicating a robust genomic structure with minimal redundancy (Figure 7). The *B. duperreyi* genome also had a low proportion of fragmented (F:1.1%) and missing (M:4.2%) orthologs. These results positioned *Bassiana duperreyi* favorably in terms of genome completeness and integrity, on par with other squamates, and highlights its potential as a reference for further genomic and evolutionary studies within this phylogenetic group. RNAseq data mappability was on average 98.42% (range 96.50-99.80%) attesting to the high quality and complete assembly of the genome.

**Figure 7.**
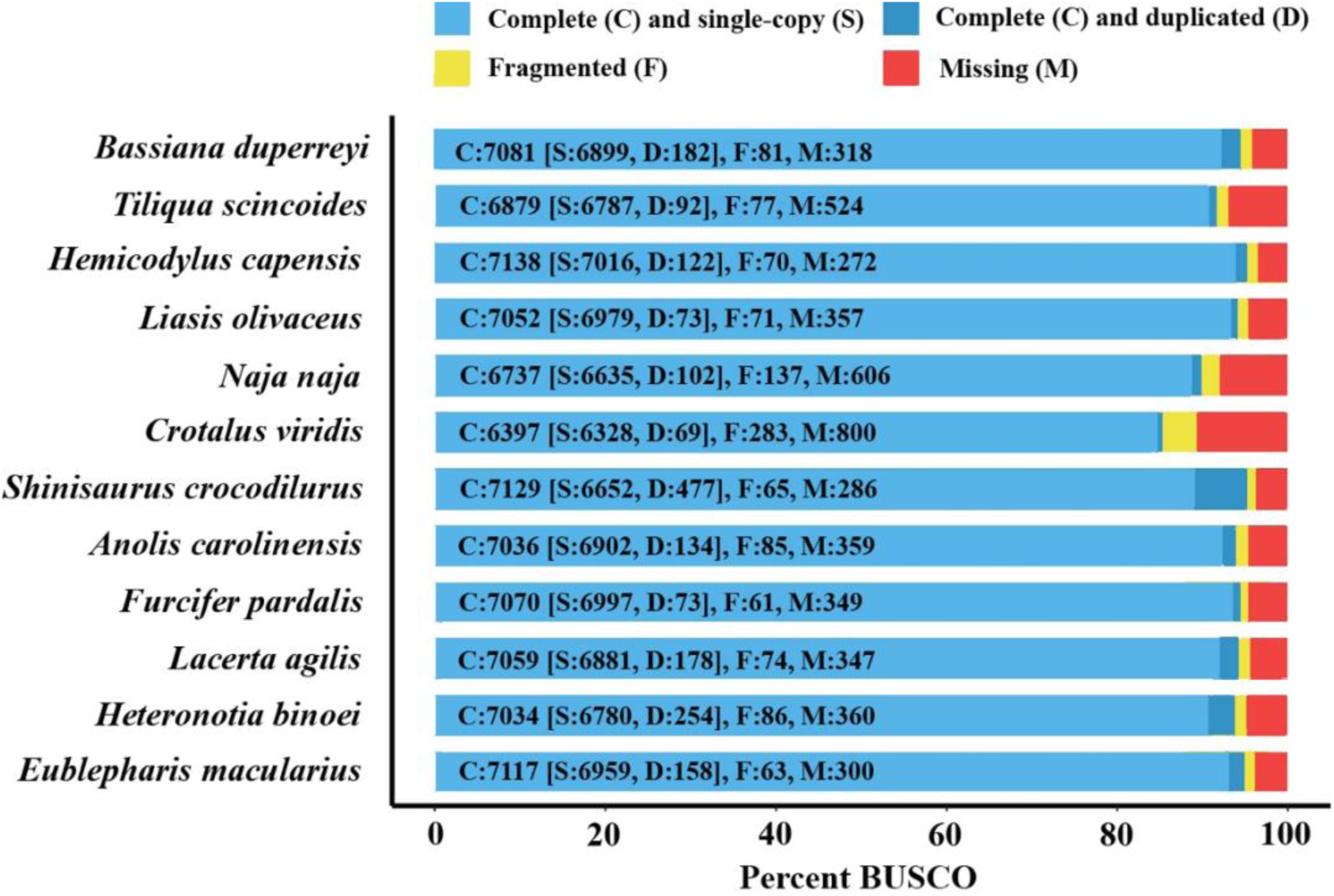
A visual representation of how complete the gene content is for each listed species genome, including *Bassiana duperreyi*, based on Benchmarking Universal Single-Copy Orthologs (BUSCO, n=7480).

### Chromosome Assembly

*Bassiana duperryii* has 2n=30 chromosomes, with seven macrochromosomes including the sex chromosomes (Figure 3). The distinction between macro and microchromosomes typically relies on a bimodal distribution of size, however other characteristics such as GC content provide additional evidence for this classification (Waters *et al*. 2021). The median GC content of 10 Kbp windows for the six largest scaffolds (representing macrochromosomes) ranged between 41.63% and 42.38%, with the X chromosome scaffold at 42.46%. In contrast, scaffolds representing chromosomes 7 and 8 had a GC content of 43.29% and 43.25%, respectively (Figure 8). The remaining six scaffolds ordered by decreasing length had a GC content of between 42.89% and 46.67% characteristic of microchromosomes in other squamates. This is consistent with the high levels of inter-chromosome contact in the HiC contact map for BASDUscf8 and other microchromosomes.

**Figure 8.**
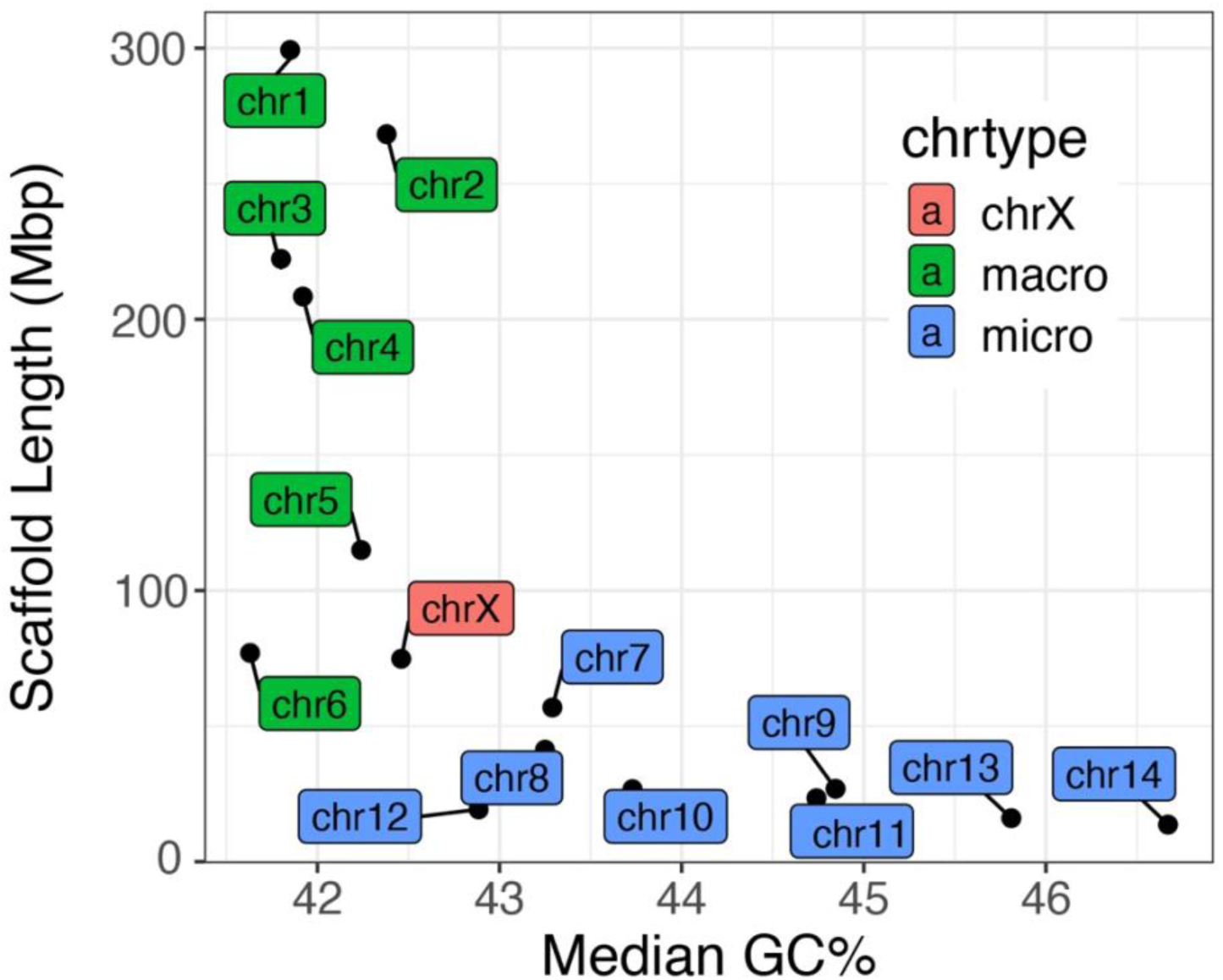
Microchromosomes are characterised by higher CG content than macrochromosomes. Median GC content in 10 Kbp windows of scaffolds vs length of scaffolds representing macrochromosomes (green), the X chromosome (red) and microchomosomes (blue).

Unlike mammals, reptiles (including most birds) show a high level of chromosomal homology across species (Waters et al. 2021). Figure 9 shows synteny conservation between *B. duperreyi* and representative squamate species. Apart from a handful of internal rearrangements, the major scaffolds of *Tiliqua scincoides* and *B. duperreyi* corresponded well, including the X chromosome (BASDUscf7.2). When compared with other genomes in the analysis, the *B. duperreyi* genome showed a high degree of evolutionary conservation with respect to both chromosomal arrangement and gene order. Our ability to recover this relationship

**Figure 9.**
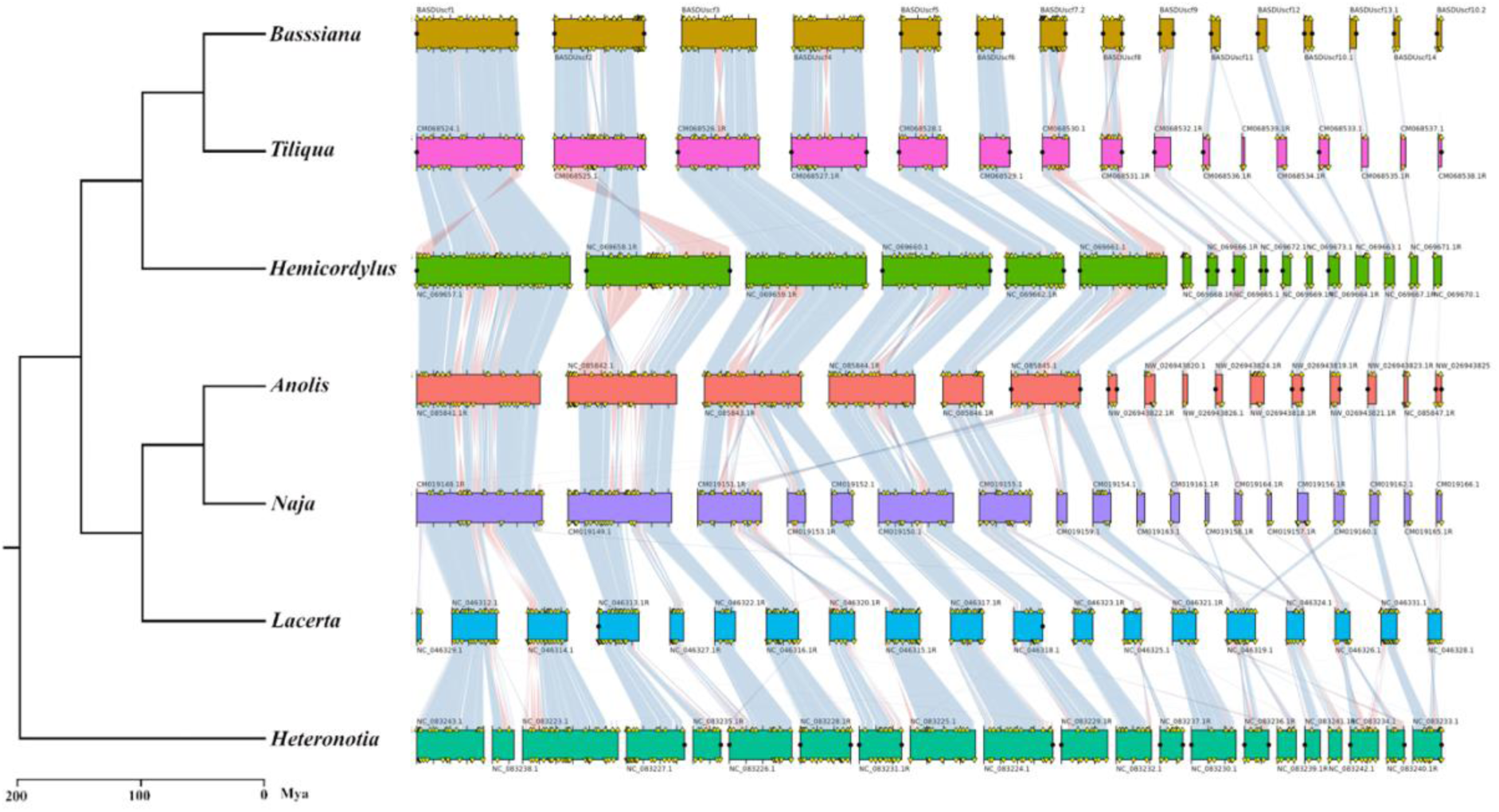
Synteny conservation of BUSCO homologs for *Bassiana duperreyi* and squamates with chromosome level assemblies including representative skink, iguanid, snake and gecko lineages (Table S7). Synteny blocks corresponding to each species are aligned horizontally, highlighting conserved chromosomal segments across the genomes. The syntenic blocks are connected by ribbons that represent homologous regions shared between species, with the varying colours denoting segments of inverted gene order. Duplicated BUSCO genes are marked with yellow triangles. Predicted telomeres are marked with black circles.

Scaffold BASDUscf7.2 of 74.8 Mbp was identified as the X chromosome based on the median read depth for 10K bp sliding windows. Read depth was half of the genome median with 17.5*x* for the PacBio HiFi, 31.8*x* for ONT and 36.3*x* for Illumina data. This putative X chromosome scaffold lacked one telomere admitting the possibility that other X chromosome sequence was present in the assembly (possibly pseudoautosomal). A total of 137 scaffolds could not be reliably mapped to a chromosome or other elements of the assembly (rDNA or centromeric satellite repeats) and were thus identified as a set containing putative Y chromosome scaffolds. We mapped Y-specific contigs (Dissanayake *et al*. 2020) to identify the Y-specific scaffolds. The assembly of the Y chromosome was fragmented with 21 scaffolds ranging in length from 56 Kbp to 6.4 Mbp and a total length of 34.5 Mb (11 ≥ 1 Mbp for a total length of 30.7 Mbp) (Figure S10). Further curation is required to improve representation of the *B. duperreyi* Y chromosome.

### Mitochondrial Genome

The *Bassiana duperreyi* mitochondrial genome was 17,506 bp in size with 37 intact genes without frameshift mutations. It consisted of 22 tRNAs, 13 protein coding genes, 2 ribosomal RNA genes and the control region (Figure S5), so was typical of the vertebrate mitochondrial genome. Base composition was A = 32.83%, C = 27.73%, G = 13.89% and T = 25.55%.

### Annotation

An estimated 53.1% (832.6 Mbp) of the *B. duperreyi* genome was composed of repetitive sequences, including interspersed repeats, small RNAs and simple and low complexity tandem repeats (Figure 10). DNA transposons were the most common repetitive element (9.26% of the genome) and are dominated by TcMar-Tigger and hAT elements. While the abundance of these elements is reported to be highly variable in squamate genomes, they make up a larger percentage of the *B. duperreyi* genome than typically found in lizards (Pasquesi *et al*. 2018). CR1, BovB and L2 elements were the dominant long interspersed elements (6.69% of the genome), which is consistent with other squamate genomes (Pasquesi *et al*. 2018). The *B. duperreyi* genome also appeared to have a significant proportion of Helitron rolling-circle (2.13%) transposable elements. More than half of all repeat content was unclassified and did not correspond to any element in the *RepeatModeler* libraries. The number of elements masked and their relative abundances are presented in the supplementary material (Table S9).

**Figure 10.**
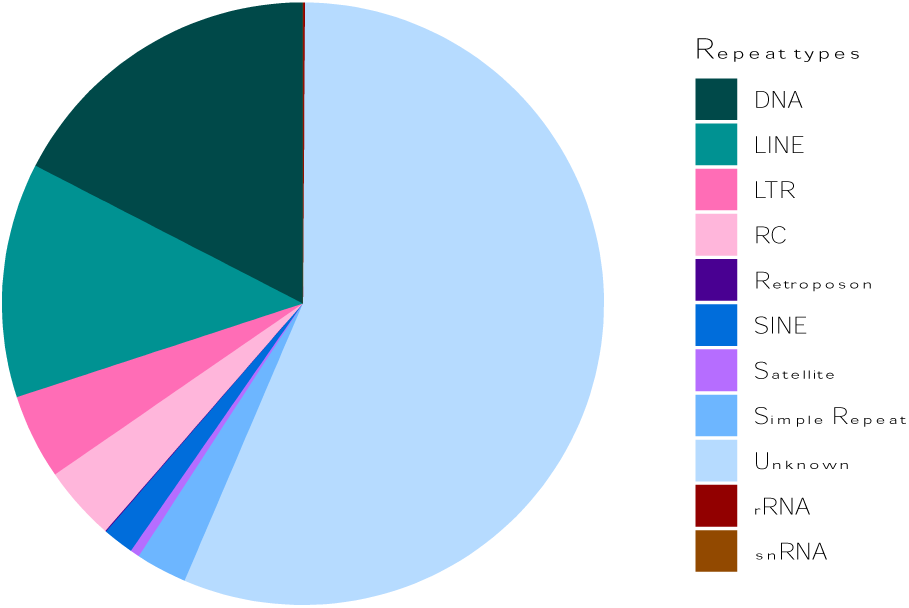
Proportion of different repeat classes in the *Bassiana duperreyi* genome. Abbreviations: DNA, DNA Transposons; LINE, Long Interspersed Nuclear Element; LTR, Long Terminal Repeat; RC Rolling Circle, mobile elements using rolling circle replication; SINE, Short Interspersed Nuclear Element; rRNA, DNA transcribed to rRNA; snRNA, DNA transcribed to snRNA [Refer to Table S10 for a detailed breakdown].

Transcriptome assembly produced 3.3 million transcripts across 35 samples (range: 50,625– 179,298, average = 95,456). A large proportion of these transcripts (range: 35.5–62.8%, average = 42.8%) aligned to the UniProt-SwissProt protein sequences, suggestive of high-quality assemblies. A total of 2,500–15,477 full length ORFs were detected for sequences aligned to the UniProt. A further 4,356–29,539 ORFs >50 amino acids with start and stop codons were detected for transcripts that did not align to UniProt. A subset of non-redundant transcripts were utilized for *de novo* gene annotations.

Genome annotation using *Augustus* predicted 19,128 genes and transcripts, of which 17,962 had a match to a Uniprot_Swissprot/Uniprot_TrEMBL protein sequence, and 17,442 were assigned a gene name. The quality of the annotation was further validated using RNAseq data from 35 samples, with an average 51.9% (ranging from 33.3% to 75.5%) of aligned reads assigned to annotated exons, indicating a reasonable level of correspondence between the predicted gene models and the observed transcriptome.

There were 13 scaffolds identified as putative rDNA scaffolds based on their alignments with 18S and 28S subunit sequences of deuterostomes. These scaffolds ranged in size between 19.1 to 347.9 Kbp. There were six small scaffolds (34.2-177.8 Kbp) that had >50% of their sequences aligning to centromeric satellite repeat (CEN187).

With respect to the sex chromosomes, we extracted and compiled a list of genes located on the X and Y chromosome scaffolds into a separate table available in the supplementary material (refer to supplementary file Table_S11.xlsx). A preliminary analysis of these gene did not reveal any obvious candidates for the master sex-determining gene. This assessment was based on both existing knowledge of sex determining genes or gene families in vertebrates, and a gene function search using Panther (https://pantherdb.org). Determining the mode of sex determination (dominance or dosage) and identifying potential master sex-determining genes on the sex chromosomes requires further investigation and is beyond the scope of this paper.

## Conclusion

Here we present a high-quality genome assembly of the Australian alpine three-lined skink *Bassiana duperreyi* (Gray 1838). The quality of the genome assembly and annotation compares well with other chromosome-length assemblies (Table S7) and is among the best for any species of Scincidae, despite the sequence data being restricted to “long” PacBio and ONT reads rather than “ultralong” reads. We have chromosome length scaffolds, each with a well-defined centromere and many telomere to telomere. The non-recombining region of the X chromososome was assembled as a single scaffold, although the pseudoautosomal region was not identified, it is likely represented among the unassembled regions or unassigned scaffolds lacking a telomeric sequence. The Y chromosome remains fragmented across multiple scaffolds. This annotated assembly for the alpine three-lined skink was generated as part of the AusARG initiative of Bioplatforms Australia, to contribute to the suite of high-quality genomes available for Australian reptiles and amphibians as a national resource. We anticipate that this reference genome will serve to accelerate comparative genomics and evolutionary research on this and other species. As an exemplar of a well studies oviparous taxon, the *B. duperreyi* reference assembly will provide a solid basis for genomic studies of the evolution of viviparity and placentation across the Scincidae (Stewart and Thompson 1996; Foster *et al*. 2022) and for studies of the genetic basis for reprogramming of sexual development under the influence of environmental temperature (Dissanayake *et al*. 2021a,b).

## Funding

This work was supported by the AusARG initiative funded by Bioplatforms Australia and the Australian Research Council (DP220101429), and National Health and Medical Research Council (APP2021172).

## Availability of supporting data

The supplementary file contains a description of all supplemental materials, which include Tables showing software used in the preparation of this paper, outcomes of the sequencing on the four sequencing platforms used, and figures in support of statements on the quality of data. The authors affirm that all other data necessary for confirming the conclusions of the article are present within the article, figures, and tables. The annotated assembly can be accessed from NCBI and all reads used in support of the assembly are lodged with the Short Read Archive. Accession numbers are provided in the main text and the Supplementary Tables (Tables S2-S6). High resolution versions of Figures and custom scripts used to conduct the analyses are at https://github.com/kango2/basdu.

## Author Contributions

All authors contributed to the writing and editing of drafts of this manuscript. In addition, A.G. was the AusARG project lead and responsible for securing the funding; A.L.M.R. contributed to the development of assembly pipelines; B.J.H. was responsible for analyses of the comparative performance of the assembly and final submission; D.O’M collected the initial samples and undertook preliminary assembly of the transcriptome and genome; D.S.B.D – collected samples and the initial conceptual work; H.R.P. led the assembly and development of related workflows and pipelines; I.W.D. provided oversight of the data generation and supervision of subsequent analysis; J.C. developed the annotation workflow and pipelines and read depth analyses; J.M.H contributed to data generation and associated quality control and submission to NCBI; K.A. was responsible under the supervision of H.R.P for data curation and management, constructing the automated assembly and annotation workflows, for the manual curation of the assembly & analysis and post-assembly analysis; P.W. with H.R.P. provided oversight of the assembly and annotation, interpretation of the X and Y scaffolds; R.J.E. provided scripts for cross species alignments and their display; T.B. took the lead on the analysis of repeat structure.

## Acknowledgements

We acknowledge the provision of computing and data resources provided by the Australian BioCommons Leadership Share (ABLeS) program. This program is co-funded by Bioplatforms Australia (enabled by NCRIS) and the National Computational Infrastructure (NCI).

## Competing interest

H.R.P., I.W.D., A.G. have previously received travel and accommodation expenses from ONT and/or PacBio to speak at conferences. I.W.D. has a paid consultant role with Sequin Pty Ltd. H.R.P. holds equity in ONT, PacBio and Illumina. The authors declare no other competing interests.

