## Supplementary figures and tables for "A genome assembly and annotation for the Australian alpine skink *Bassiana duperreyi* using long-read technologies"

### **List of Tables**

**Table S1.** A list of software used for the analyses reported in this paper.

**Table S2.** Summary statistics for the raw Illumina DNA sequence data.

##### **Table S3.** Summary statistics for the raw PacBio HiFi sequence data used for the assembly.

##### **Table S4.** Summary statistics for the raw Oxford Nanopore sequence data used for the assembly.

##### **Table S5.** Summary statistics for the HiC sequence data used to scaffold the assembly.

**Table S6.** Summary statistics for the raw Illumina RNA sequence data used to assemble the transcriptome and for annotation.

**Table S7.** Comparison of the *Bassiana duperreyi* assembly with selected other chromosome level assemblies for chicken and squamates.

**Table S8.** Centromeric motifs from the genome assembly of *Bassiana duperreyi*.

**Table S9.** Length of centromeric repeats on the major scaffolds of the genome assembly for *Bassiana duperreyi*.

**Table S10.** Summary of the copy number and percentage of the *Bassiana duperreyi* genome covered by repeats.

**Table S11.** List of genes annotated on the X and Y chromosomes of *Bassiana duperreyi*. Refer separate file Table_S11.xlsx.

**Table S12.** List of scaffolds in the assembly of *Bassiana duperreyi* and their assignment where possible to chromosomes. Refer separate file Table_12.xlsx

### **List of Figures**

**Figure S1.** Comparison of average read quality values versus read length for the two sequencing technologies: Oxford Nanopore Technologies and PacBio HiFi.

**Figure S2**. Manual breaks arising from internal telomeric sequence.

**Figure S3.** Mapping of Y-enriched k-mer contigs to the assembly. A high resolution version of this figure can be found at <https://github.com/kango2/basdu>.

**Figure S4.** A plot of the 15 longest scaffolds (corresponding to the number of chromosomes of *Bassiana duperreyi*) for the YaHS assembly. A high-resolution version of this figure can be found at <https://github.com/kango2/basdu>.

**Figure S5**. Annotation of the mitochondrial genome of *Bassiana duperreyi* using mitoHiFi.

**Figure S6.** HiC contact maps displayed using Juicer for the curated assembly.

### **Custom Scripts** (available from <https://github.com/kango2/basdu>).

1. **pacbiobam2fastx.sh** – A custom script to remove any reads containing PacBio adapter sequences and convert the .bam files to FASTQ.
2. **calculateGC.py** – A custom script to calculate GC content in non-overlapping sliding windows of 10 Kbp.
3. **blastxtranslation.pl** – A custom script to obtain putative open reading frames and corresponding amino acid sequences
4. **processtrftelo.py** – A custom script to identify regions >600 bp that contained conserved vertebrate telomeric repeat motif (TTAGGG).

**Table S1.** A list of software and databases used for the analyses reported in this paper. Included are the use to which the software or database was put, the version number used, the source of the latest release of the software or database, and the associated published reference if applicable.

| **Software** | **Use case** | **Version** | **URL (latest release)** | **Reference** |
| --- | --- | --- | --- | --- |
| *AGAT* | Conversion from bam to gff3 | 1.4.0 | https://github.com/NBISweden/AGAT | – |
| *agptools* | Splitting scaffolds | – | https://github.com/WarrenLab/agptools | – |
| *Arima Genomics alignment pipeline* | Aligning HiC data | – | https://github.com/ArimaGenomics/mapping_pipeline | – |
| *Augustus* | De novo gene annotations | 3.4.0 | https://github.com/Gaius-Augustus/Augustus | Stanke *et al.* 2008 |
| *BLAST* | Finds regions of local similarity between sequences | 2.14.1 | https://blast.ncbi.nlm.nih.gov/Blast.cgi | - |
| *BUSCO* | Assess completeness of conserved genes | 5.4.7 | https://gitlab.com/ezlab/busco | Manni *et al.* 2021 |
| *buttery-eel* | ONT basecalling | 0.4.2 | https://github.com/Psy-Fer/buttery-eel | Samarakoon *et al.* 2023a |
| *bwa-mem2* | Short-read alignments | 2.2.1 | https://github.com/bwa-mem2/bwa-mem2 | Vasimuddin *et al.* 2019 |
| CD-HIT | Clustering of redundant transcript sequences across multiple samples | 4.8.1 | https://sites.google.com/view/cd-hit | Fu *et al*. 2012 |
| Chromosyn | Create BUSCO synteny plots | 1.3.0 | https://github.com/slimsuite/chromsyn | Edwards *et al.* 2022 |
| Cutadapt | Removing adaptor sequence | 3.7 | https://github.com/marcelm/cutadapt | Martin 2011 |
| DeepConsensus | Basecalling from subreads | 1.2.0 | https://github.com/google/deepconsensus | Baid *et al.* 2023 |
| diamond | Aligning transcriptomic and peptide sequences to uniprot databases for annotation | 2.1.9 | https://github.com/bbuchfink/diamond | Buchfink *et al.* 2021 |
| dorado | ONT basecalling | 7.2.13 | https://github.com/nanoporetech/dorado | – |
| genometools | Parse gff3 annotation files | 1.6.2 | https://github.com/genometools/genometools | Gremme *et al.* 2013 |
| hifiasm | Assembly construction | 0.19.8 | https://github.com/chhylp123/hifiasm | Cheng *et al.* 2021a, 2022 |
| Inspector | Evaluating long read de-novo assembly results | 1.2 | https://github.com/Maggi-Chen/Inspector | Cheng *et al.* 2021b |
| Merqury | Evaluate assembly completeness with kmers | 1.3 | https://github.com/marbl/merqury | Rhie *et al.* 2020 |
| Meryl | Generate kmer database | 1.4.1 | https://github.com/marbl/meryl | Rhie *et al.* 2020 |
| MitoHiFi | Mitochondrial genome annotation | 3.2.2 | https://github.com/marcelauliano/MitoHiFi | Uliano-Silva *et al.* 2023 |
| minimap2 | Long-read alignments, Alignment for gene model training and prediction | 2.17 (assembly), 2.26 (annotation) | https://github.com/lh3/minimap2 | Li 2018 |
| RepeatMasker | Repeat annotations | 4.1.2-p1 | https://github.com/rmhubley/RepeatMasker | Smit *et al.* 2013-2015 |
| RepeatModeler | Repeat annotations | 2.0.4 | https://github.com/Dfam-consortium/RepeatModeler | Smit *et al.* 2008-2015 |
| Samtools | SAM/BAM viewing, manipulation and calculations | 1.19 (assembly),1.19.2 (annotatoion) | <https://github.com/samtools/samtools> | Danecek *et al.* 2021 |
| Subread | Alignment of RNAseq (subread-align) | 2.0.6 | https://subread.sourceforge.net | Liao *et al.* 2013 |
| slow5tools | Conversion of ONT file format | 1.1.0 | https://github.com/hasindu2008/slow5tools | Samarakoon *et al.* 2023b |
| TRASH | Centromeric satellite repeat annotations | 1.12 | https://github.com/vlothec/TRASH | Wlodzimierz *et al.* 2023 |
| TRF | Tandem repeat annotations including telomeres | 4.09.1 | https://github.com/Benson-Genomics-Lab/TRF | Benson 1999 |
| Trimmomatic | Trimming of illumina sequence data | 0.39 | https://github.com/usadellab/Trimmomatic | Bolger *et al.* 2014 |
| Trinity | Transcriptome assembly | 2.12.0 | https://github.com/trinityrnaseq/trinityrnaseq | Grabherr *et al.* 2011 |
| VecScreen and UniVec database | Screen for vector sequence contamination | – | https://www.ncbi.nlm.nih.gov/tools/vecscreen | – |
| YaHS | Scaffolding with HiC | 1.1 | https://github.com/c-zhou/YaHS | Zhou *et al.* 2022 |

**Table S2.** Summary statistics for the raw Illumina DNA sequence data. Values separated by a semicolon are for the first and second member of the paired end reads.

| SpecimenID (UC<Aus>) | Tissue | LibraryID | SRA | No. of Bases | No. of Reads | Mean Read Length | No. of missing values (Ns) |
| --- | --- | --- | --- | --- | --- | --- | --- |
| DDBD_364 | Muscle | 350747_L001 | SRR25773554 | 27792097427;27903978856 | 115441574;115441574 | 240.7;241.7 | 25344305;2347537 |
| DDBD_364 | Muscle | 350747_L002 | SRR25773554 | 27405045539;27511746903 | 113877370;113877370 | 240.7;241.6 | 23914990;1583932 |
| Bd64 | Brain | bd64.180bp | SRR12196015 | 9670750200;9670750200 | 96707502;96707502 | 100;100 |  |
| Bd65 | Brain | bd65.180bp | SRR12196012 | 8141146800;8141146800 | 81411468;81411468 | 100;100 |  |

##### **Table S3.** Summary statistics for the raw PacBio HiFi sequence data used for the assembly.

| SpecimenID (UC<Aus>) | Tissue | Flow Cell | SRA | No. of Bases | No. of Reads | Mean Read Length | N50 | N90 |
| --- | --- | --- | --- | --- | --- | --- | --- | --- |
| DDBD_364 | Blood | DA060219 | SRR28919959 | 26008467217 | 1690986 | 15381 | 15632 | 12052 |
| DDBD_364 | Blood | DA060220 | SRR28919959 | 26428916467 | 1704390 | 15506 | 15774 | 12156 |

##### **Table S4.** Summary statistics for the raw Oxford Nanopore sequence data used for the assembly.

| SpecimenID (UC<Aus>) | Tissue | Flow cell | SRA | No. of Bases | No. of Reads | Mean Read Length | N50 | N90 |
| --- | --- | --- | --- | --- | --- | --- | --- | --- |
| DDBD_364 | Muscle | PAG18312 | SRR25773548 | 37261710121 | 7614955 | 4,893 | 11181 | 2096 |
| DDBD_364 | Muscle | PAG18256 | SRR25773549 | 67210354449 | 14429383 | 4,658 | 10812 | 1964 |

##### **Table S5.** Summary statistics for the HiC sequence data used to scaffold the assembly. Values separated by a semicolon are for the first and second member of the paired end reads.

| SpecimenID (UC<Aus>) | Tissue | Library | SRA | No. of Bases | No. of Reads | Mean Read Length | No. of missing values (Ns) |
| --- | --- | --- | --- | --- | --- | --- | --- |
| DDBD_364 | Liver | 350769_L001 | SRR25773553 | 20582513889;20582513889 | 136308039;136308039 | 151.0;151.0 | 100216:173307 |
| DDBD_364 | Liver | 350769_L002 | SRR25773553 | 20329523053;20329523053 | 134632603;134632603 | 151.0;151.0 | 102601:112369 |

| **SpecimenID (UC<Aus>)** | **Tissue** | **LibraryID** | **SRA** | **No. of Bases** | **No. of Reads** | **Mean Read Length** | **No. of ambiguous bases (Ns)** | **Genes** | **Trans-cripts** | **Uniprot** | **Un-aligned** | **orf50 plus** | **Aln full orf** | **Unaln full orf50plus** |
| --- | --- | --- | --- | --- | --- | --- | --- | --- | --- | --- | --- | --- | --- | --- |
| DDBD_532 | Brain | 350723 | SRR25773540 | 3446462387; 3445697637 | 45657586; 45657586 | 75.5; 75.5 | 92960; 158654 | 83283 | 114620 | 43939 | 70681 | 90851 | 3514 | 14636 |
| DDBD_532 | Heart | 350724 | SRR25773557 | 3380233266; 3379602666 | 44772671; 44772671 | 75.5; 75.5 | 82626; 149362 | 64692 | 85668 | 32802 | 52866 | 68218 | 2500 | 10664 |
| DDBD_141 | Testes | 350725 | SRR28919964 | 3489436749; 3484700814 | 46197880; 46197880 | 75.5; 75.4 | 100485; 164225 | 72334 | 104805 | 41081 | 63724 | 84606 | 6503 | 11253 |
| DDBD_533 | Brain | 350726 | SRR25773556 | 3559742707; 3558803028 | 47151887; 47151887 | 75.5; 75.5 | 92540; 161518 | 84125 | 116707 | 44736 | 71971 | 92783 | 4735 | 15149 |
| DDBD_533 | Heart | 350727 | SRR25773555 | 3722825550; 3722016369 | 49302100; 49302100 | 75.5; 75.5 | 92506; 161538 | 73355 | 100369 | 37732 | 62637 | 79778 | 3118 | 13216 |
| DDBD_533 | Ovary | 350728 | SRR28919963 | 3937818340; 3933721019 | 52140064; 52140064 | 75.5; 75.5 | 186719; 243960 | 34021 | 50625 | 30692 | 19933 | 45398 | 8813 | 4356 |
| DDBD_358 | Brain | V33 | [SRR25773559](https://dataview.ncbi.nlm.nih.gov/object/SRR25773559) | 2418450300 | 32246004 | 75 | 51848 | 62213 | 75968 | 30990 | 44978 | 61487 | 3425 | 10521 |
| DDBD_397 | Brain | V35 | [SRR25773558](https://dataview.ncbi.nlm.nih.gov/object/SRR25773558) | 2526695325 | 33689271 | 75 | 53230 | 68317 | 84160 | 33627 | 50533 | 67635 | 3415 | 11558 |
| DDBD_272 | Brain | V36 | [SRR25773547](https://dataview.ncbi.nlm.nih.gov/object/SRR25773547) | 2368019175 | 31573589 | 75 | 49661 | 62579 | 75614 | 31022 | 44592 | 61291 | 3161 | 10474 |
| DDBD_275 | Brain | V37 | [SRR25773546](https://dataview.ncbi.nlm.nih.gov/object/SRR25773546) | 2165833425 | 28877779 | 75 | 46785 | 60935 | 72801 | 29833 | 42968 | 59119 | 3241 | 9716 |
| DDBD_329 | Brain | V38 | SRR25773545 | 2437856550 | 32504754 | 75 | 52490 | 60199 | 73269 | 30304 | 42965 | 59545 | 3250 | 10315 |
| DDBD_322 | Brain | V40 | [SRR25773544](https://dataview.ncbi.nlm.nih.gov/object/SRR25773544) | 2260340850 | 30137878 | 75 | 48796 | 61652 | 75118 | 30653 | 44465 | 60736 | 2622 | 10607 |
| DDBD_218 | Brain | V41 | [SRR25773543](https://dataview.ncbi.nlm.nih.gov/object/SRR25773543) | 2293725150 | 30583002 | 75 | 48393 | 61791 | 74131 | 30613 | 43518 | 59973 | 3233 | 9756 |
| DDBD_400 | Brain | V43 | [SRR25773542](https://dataview.ncbi.nlm.nih.gov/object/SRR25773542) | 2621211525 | 34949487 | 75 | 55374 | 65268 | 80789 | 32384 | 48405 | 65182 | 3417 | 11196 |
| DDBD_220 | Brain | V44 | [SRR25773541](https://dataview.ncbi.nlm.nih.gov/object/SRR25773541) | 2679905400 | 35732072 | 75 | 56491 | 64517 | 79819 | 32435 | 47384 | 64842 | 3972 | 11221 |
| DOM_Bd59 | Brain | DOM_3 | SRR28919958 | 2730432485; 2730432485 | 27033985; 27033985 | 101; 101 | 4490742;  2512445 | 71795 | 117842 | 49324 | 68518 | 98372 | 12148 | 19462 |
| DOM_Bd13 | Brain | DOM_4 | SRR28919957 | 3374000950; 3374000950 | 33405950; 33405950 | 101; 101 | 5563608;  3062232 | 74248 | 122994 | 50600 | 72394 | 102754 | 12798 | 20847 |
| DOM_Bd65 | Brain | DOM_5 | SRR28919956 | 3115022406; 3115022406 | 30841806; 30841806 | 101; 101 | 5126631;  2840125 | 66182 | 107248 | 46081 | 61167 | 89992 | 11566 | 17423 |
| DOM_Bd59 | Testis | DOM_6_7_8 | SRR28919955 |  |  |  |  |  |  |  |  |  |  |  |
| DOM_Bd13 | Testis | DOM_6_7_8 | SRR28919955 | 3745492484; 3745492484 | 37084084; 37084084 | 101; 101 | 6150098;  3427706 | 98206 | 179298 | 63635 | 115663 | 148262 | 15477 | 29539 |
| DOM_AA045727 | Testis | DOM_6_7_8 | SRR28919955 |  |  |  |  |  |  |  |  |  |  |  |
| DOM_KNP | Brain | DOM_9 | SRR28919954 | 1582227014; 1582227014 | 15665614; 15665614 | 101; 101 | 2607055;  1457795 | 56442 | 86596 | 38560 | 48036 | 72731 | 9007 | 13607 |
| DOM_Bd64 | Brain | DOM_10 | SRR28919953 | 2071723009; 2071723009 | 20512109; 20512109 | 101; 101 | 3412857;  1899211 | 54489 | 80613 | 36554 | 44059 | 68103 | 7993 | 12480 |
| DOM_AA045725 | Brain | DOM_11 | SRR28919952 | 2215957170; 2215957170 | 21940170; 21940170 | 101; 101 | 3659757;  2025331 | 63193 | 101236 | 43932 | 57304 | 85048 | 11065 | 16308 |
| DOM_AA045726 | Ovary | DOM_12_13_14 | SRR28919962 |  |  |  |  |  |  |  |  |  |  |  |
| DOM_KNP | Ovary | DOM_12_13_14 | SRR28919962 | 2148252426; 2148252426 | 21269826; 21269826 | 101; 101 | 3527790;  1972882 | 31392 | 54332 | 34139 | 20193 | 49499 | 13391 | 5552 |
| DOM_AA045725 | Ovary | DOM_12_13_14 | SRR28919962 |  |  |  |  |  |  |  |  |  |  |  |
| DOM_Bd13 | Liver | DOM_15 | SRR28919961 | 2066009843; 2066009843 | 20455543; 20455543 | 101; 101 | 3398544;  1879237 | 47646 | 68718 | 29010 | 39708 | 57259 | 6653 | 10434 |
| DOM_KNP | Liver | DOM_16 | SRR28919960 | 3028867689; 3028867689 | 29988789; 29988789 | 101; 101 | 4983618;  2780107 | 54115 | 84309 | 35885 | 48424 | 70149 | 8768 | 13242 |
| BD2001_ONG | Uterus | BD2001_ONG | SRR12656980 | 4075826822; 4074182331 | 40564798; 40564798 | 100.48; 100.44 | 182416; 386369 | 89647 | 151504 | 60655 | 90849 | 125612 | 14124 | 23448 |
| BD1815_ONG | Uterus | BD1815_ONG | SRR12656981 | 3911308573; 3909429457 | 38926260; 38926260 | 100.48; 100.44 | 70617; 373741 | 75772 | 128240 | 54479 | 73761 | 107949 | 13717 | 19780 |
| BD1813_ONG | Uterus | BD1813_ONG | SRR12656982 | 4431864293; 4430107467 | 44123132; 44123132 | 100.44; 100.40 | 133481; 421290 | 73872 | 124585 | 53007 | 71578 | 104980 | 13976 | 19307 |
| BD1812_ONG | Uterus | BD1812_ONG | SRR12656983 | 3668468719; 3667130706 | 36532202; 36532202 | 100.41; 100.38 | 177531; 347478 | 75558 | 127520 | 53802 | 73718 | 107650 | 12505 | 20413 |
| BD1811_ONG | Uterus | BD1811_ONG | SRR12656984 | 3295799239; 3294840594 | 32817258; 32817258 | 100.43; 100.40 | 193639; 294801 | 76208 | 127756 | 53437 | 74319 | 107513 | 12092 | 20130 |
| BD1809_OLG | Uterus | BD1809_OLG | SRR12656989 | 3165918687; 3166231370 | 31534074; 31534074 | 100.40; 100.40 | 185572; 303598 | 46427 | 67059 | 30704 | 36355 | 57239 | 7839 | 9907 |
| BD1806_OLG | Uterus | BD1806_OLG | SRR12657000 | 3945760567; 3945526459 | 39300685; 39300685 | 100.40; 100.4 | 531860; 375717 | 55166 | 89470 | 40746 | 48724 | 76615 | 10527 | 14426 |
| BD1805_OLG | Uterus | BD1805_OLG | SRR12657011 | 3206362176; 3206659458 | 31926933; 31926933 | 100.42;100.40 | 93928; 310531 | 50541 | 75421 | 33708 | 41713 | 64055 | 7523 | 11360 |
| BD1802_OLG | Uterus | BD1802_OLG | SRR12657022 | 3841362849; 3840904039 | 38256768; 38256768 | 100.41; 100.3 | 406071; 361574 | 56106 | 93182 | 43166 | 50016 | 79615 | 9986 | 14119 |
| BD1801_OLG | Uterus | BD1801_OLG | SRR12657023 | 3772381731; 3772886001 | 37609872; 37609872 | 100.30; 100.3 | 388976; 369186 | 57397 | 88560 | 38359 | 50201 | 74768 | 9270 | 13224 |

**Table S7.** Comparison of the *Bassiana duperreyi* assembly with selected other chromosome level assemblies for chicken and squamates.

| Assembly | *Bassiana duperreyi* | *Anolis carolinensis* | *Gallus gallus* | *Lacerta agilis* | *Naja Naja* | *Shinisaurus crocodilurus* |
| --- | --- | --- | --- | --- | --- | --- |
| GenBank Accession | ***To be provided on acceptance*** | GCA_000090745.2 | GCA_000002315.5 | GCA_009819535.1 | GCA_009733165.1 | GCA_021292165.1 |
| Year of release | 2024 | 2014 | 2018 | 2019 | 2021 | 2021 |
| Scaffolds | 172 | 6,457 | 463 | 28 | 1,897 | 1553 |
| Total Length | 1.57 Gb | 1.80 Gb | 1.07 Gb | 1.39 Gb | 1.77 Gb | 2.19 Gb |
| Scaffold N50 | 222,269,761 | 150,641,568 | 91,315,245 | 86,565,987 | 224,088,896 | 296,945,371 |
| Scaffold N90 | 26,766,351 | 408,349 | 10,762,512 | 43,910,048 | 23,367,680 | 10,640,786 |
| Scaffold L50 | 3 | 5 | 4 | 7 | 3 | 4 |
| Scaffold L90 | 11 | 416 | 19 | 16 | 14 | 16 |
| Largest contig | 299,325,919 | 263,920,458 | 197,608,386 | 133,750,839 | 375,026,955 | 321,223,554 |
| Assembly | *Hemicordylus capensis* | *Cryptoblepharus egeriae* | *Furcifer pardalis* | *Tiliqua scincoides* | *Eublepharis macularius* | *Liasis olivaceus* |
| GenBank Accession | GCA_027244095.1 | GCA_030015325.1 | GCA_030440675.1 | GCA_035046505.1 | GCA_028583425.1 | GCA_030867105.1 |
| Year of release | 2022 | 2023 | 2023 | 2024 | 2023 | 2023 |
| Scaffolds | 44 | 74 | 11 | 350 | 75 | 43 |
| Total Length | 2.29 Gb | 1.41 Gb | 1.61 Gb | 1.62 Gb | 2.24 Gb | 1.49 Gb |
| Scaffold N50 | 359,646,233 | 109,130,952 | 184,669,281 | 242,160,784 | 145,573,841 | 203,662,644 |
| Scaffold N90 | 33,073,465 | 30,261,696 | 111,427,539 | 48,696,318 | 59,511,070 | 26,295,960 |
| Scaffold L50 | 3 | 4 | 4 | 3 | 6 | 3 |
| Scaffold L90 | 8 | 12 | 8 | 9 | 15 | 10 |
| Largest contig | 459,129,262 | 269,273,026 | 317,477,549 | 314,321,002 | 255,159,772 | 332,077,321 |

**Table S8.** Centromeric motifs from the genome assembly of *Bassiana duperreyi*.

| Name | Length | Sequence |
| --- | --- | --- |
| CEN199 | 199 | AATGTTAGCCTGCATTGGGGCCAATGGTGACTCCAGTTGGAAAATGACTTCAGTTCATCTTGGTTGTCCACTCTTCTAGTTACATGAATCAAACCAGTTATTGGCACCACAGTGCTTAGATTGGTCTGCTGCATGTGTGGAAGCACTTAATTTCGTGACTTTCCAAAGAGTCACCAACTTACATGGAAGATGAAAACCA |
| CEN187 | 187 | CCTGAGTGAATTTGGGGGACGTGGTGCTAATAAATACAATTCTGAGTCAATTTGGGGTACAAGTATTTTTGGAAAAATACCAAAGGGGTACCCCTTGGTGAAAATCGCGTTTTGGGGTGCATCCAAAAAACACTGCAGACACCTTGGAAATGTAAGTGGGGGTGCCCCTGATGCTAAAAAACACAAT |

**Table S9.** Length of centromeric repeats on the major scaffolds of the genome assembly for *Bassiana duperreyi*. There are two classes of repeats, one based on a motif of 199 bp, the other on a motif of 187 bp (Table S8). N provides counts of the repeat motifs; bp provides the length of the repetitive element in bp.

| **Scaffold** | **Putative** | **CEN199** | | **CEN187** | |
| --- | --- | --- | --- | --- | --- |
|  | **Chromosome** | **N** | **bp** | **N** | **bp** |
| BASDUscf1 | chr1 | 4015 | 789999 | 0 | -- |
| BASDUscf2 | chr2 | 5447 | 1083999 | 6984 | 1305995 |
| BASDUscf3 | chr3 | 4095 | 814937 | 0 | -- |
| BASDUscf4 | chr4 | 4502 | 895997 | 663 | 123999 |
| BASDUscf5 | chr5 | 3133 | 623526 | 920 | 171999 |
| BASDUscf6 | chr6 | 3749 | 745998 | 0 | -- |
| BASDUscf7.2 | chrX | 5131 | 1020999 | 0 | -- |
| BASDUscf8 | chr7 | 3030 | 602999 | 1401 | 261998 |
| BASDUscf9 | chr8 | 3352 | 666997 | 5235 | 978993 |
| BASDUscf11 | chr9 | 3226 | 641999 | 722 | 134999 |
| BASDUscf12 | chr10 | 2317 | 460998 | 626 | 116999 |
| BASDUscf10.1 | chr11 | 2427 | 482999 | 0 | -- |
| BASDUscf13.1 | chr12 | 5819 | 1157998 | 2262 | 422997 |
| BASDUscf14 | chr13 | 945 | 187999 | 3102 | 579995 |
| BASDUscf10.2 | chr14 | 3910 | 777996 | 9726 | 1818738 |

**Table S10.** Summary of the copy number and percentage of the *Bassiana duperreyi* genome covered by repeats.

| **Class** | **Subclass** | **Counts** | **Length Masked (bp)** | **Percent of Sequence** |
| --- | --- | --- | --- | --- |
| **DNA Transposons** | | **826310** | **144277575** | **9.20** |
|  | CMC-Chapaev-3 | 8685 | 931404 | 0.06 |
|  | CMC-EnSpm | 842 | 45125 | 0.00 |
|  | Maverick | 4001 | 975298 | 0.06 |
|  | PiggyBac | 1357 | 270710 | 0.02 |
|  | Sola-1 | 897 | 134050 | 0.01 |
|  | Sola-2 | 712 | 206592 | 0.01 |
|  | TcMar | 6592 | 954065 | 0.06 |
|  | TcMar-Mariner | 3266 | 1735445 | 0.11 |
|  | TcMar-Pogo | 784 | 99785 | 0.01 |
|  | TcMar-Tc1 | 3376 | 636421 | 0.04 |
|  | TcMar-Tc2 | 66782 | 11250163 | 0.72 |
|  | TcMar-Tigger | 255551 | 61474742 | 3.92 |
|  | Zisupton | 1073 | 195438 | 0.01 |
|  | hAT | 27322 | 7069035 | 0.45 |
|  | hAT-Ac | 75546 | 10108526 | 0.64 |
|  | hAT-Blackjack | 1159 | 300318 | 0.02 |
|  | hAT-Charlie | 312887 | 38723344 | 2.47 |
|  | hAT-Pegasus | 415 | 112680 | 0.01 |
|  | hAT-Tag1 | 95 | 29983 | 0.00 |
|  | hAT-Tip100 | 54968 | 9024451 | 0.57 |
| **Retroelements** | |  |  |  |
|  | **LINE** | **366499** | **104873641** | **6.68** |
|  | CR1 | 168310 | 40579332 | 2.58 |
|  | Dong-R4 | 17523 | 7160730 | 0.46 |
|  | I | 193 | 75669 | 0.00 |
|  | I-Jockey | 16821 | 3317428 | 0.21 |
|  | L1 | 6105 | 1834221 | 0.12 |
|  | L2 | 34288 | 9971294 | 0.64 |
|  | Penelope | 3079 | 715042 | 0.05 |
|  | R2-NeSL | 267 | 139007 | 0.01 |
|  | RTE | 820 | 173071 | 0.01 |
|  | RTE-BovB | 81468 | 29552436 | 1.88 |
|  | RTE-X | 31896 | 9323190 | 0.59 |
|  | Rex-Babar | 5178 | 1988553 | 0.13 |
|  | Unassigned | 551 | 43668 | 0.00 |
|  | **LTR elements** | **80382** | **38096042** | **2.43** |
|  | Copia | 3425 | 2110967 | 0.13 |
|  | DIRS | 23613 | 13063630 | 0.83 |
|  | ERV1 | 18342 | 12842648 | 0.82 |
|  | ERVK | 2142 | 658156 | 0.04 |
|  | ERVL-MaLR | 194 | 33875 | 0.00 |
|  | Gypsy | 26785 | 6167183 | 0.39 |
|  | Ngaro | 5881 | 3219583 | 0.21 |
|  | **Helitron RC** | **95288** | **33336861** | **2.12** |
|  | **SNO Retroposon** | **3326** | **430585** | **0.03** |
|  | **SINE** | **96633** | **13831019** | **0.88** |
|  | 5S | 346 | 30302 | 0.00 |
|  | 5S-Deu-L2 | 1766 | 168819 | 0.01 |
|  | ID | 15878 | 1564437 | 0.10 |
|  | MIR | 57493 | 7637365 | 0.49 |
|  | U | 747 | 131141 | 0.01 |
|  | tRNA | 1201 | 80521 | 0.01 |
|  | tRNA-RTE | 19202 | 4218434 | 0.27 |
|  | **Total Interspersed Repeats** | | **334845723** | **21.33** |
|  | **Other** | **3047788** | **496872473** | **31.65** |
|  | Satellite | 4360 | 4194864 | 0.27 |
|  | Simple Repeat | 380215 | 22900194 | 1.46 |
|  | rRNA | 707 | 823469 | 0.05 |
|  | snRNA | 226 | 39915 | 0.00 |
|  | Unknown | 2662280 | 468914031 | 29.87 |
|  | **Total Masked** |  | **831718196** | **52.98** |

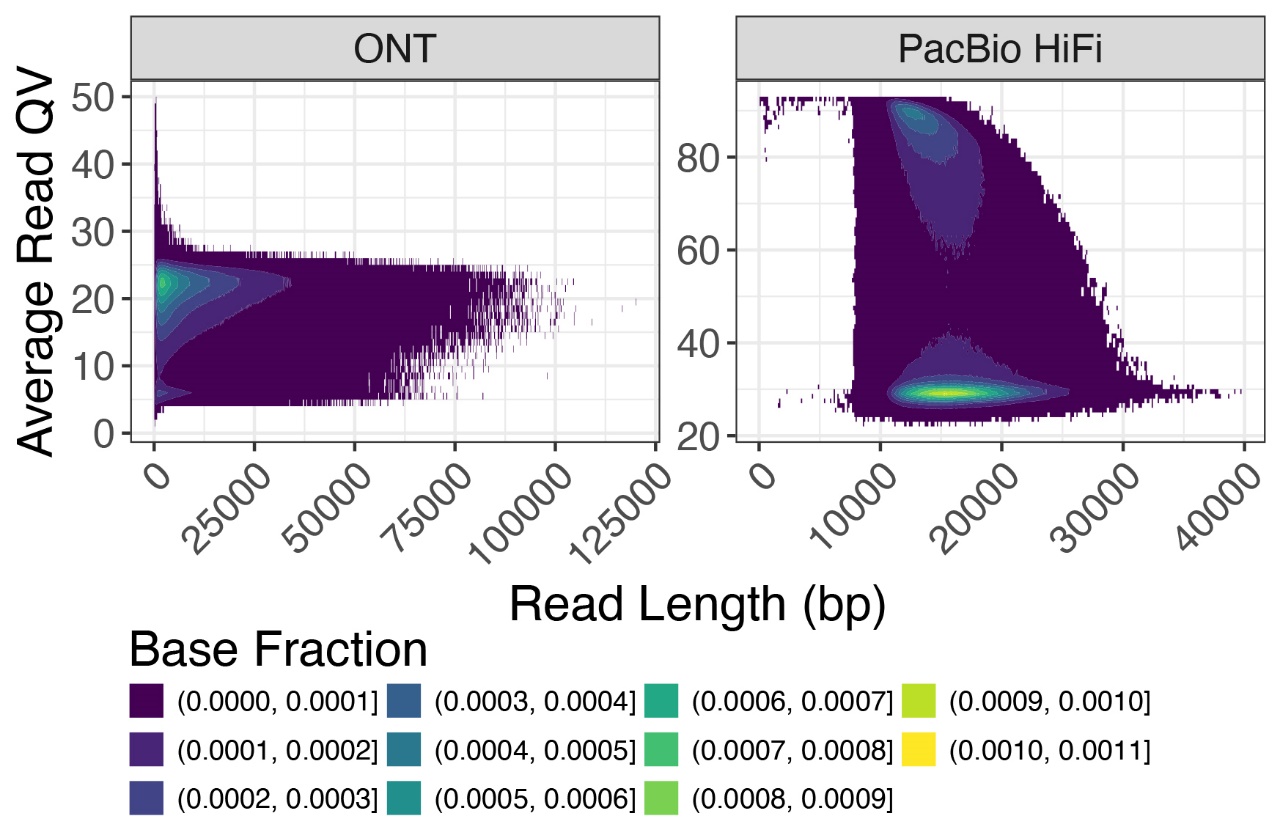

**Figure S1.** Comparison of average read quality values (QV) versus read length for the two sequencing technologies: Oxford Nanopore Technologies (ONT) and PacBio HiFi. Color intensity represents the base fraction in specified ranges, with darker colors indicating lower fractions and lighter colors indicating higher fractions. ONT reads show a broader distribution of read lengths with moderate quality values, whereas PacBio HiFi reads exhibit higher quality values with more concentrated read lengths.

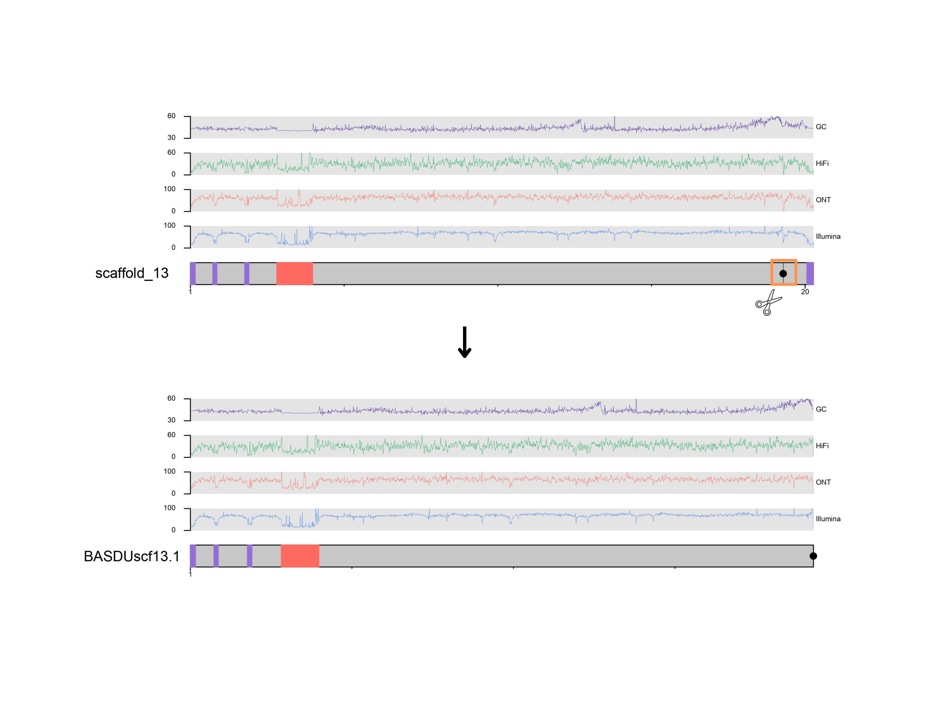

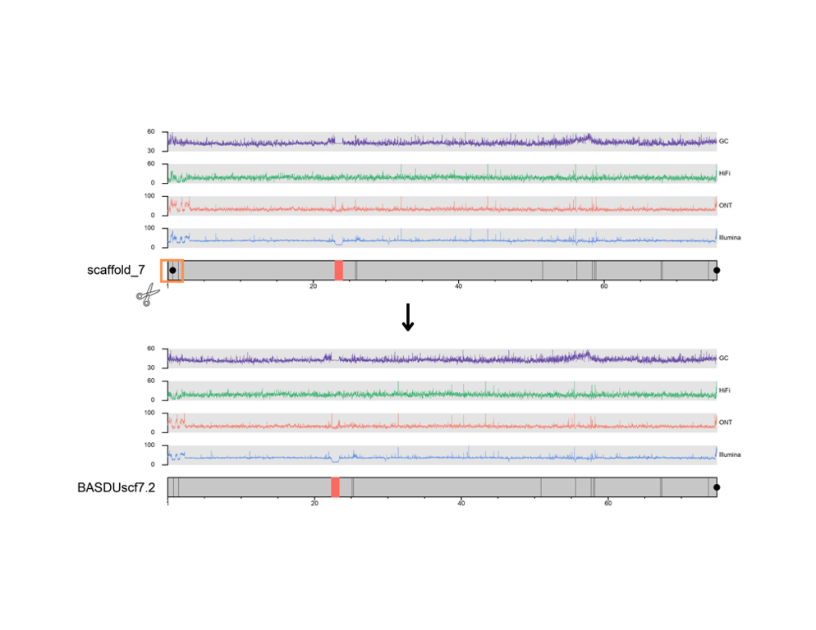

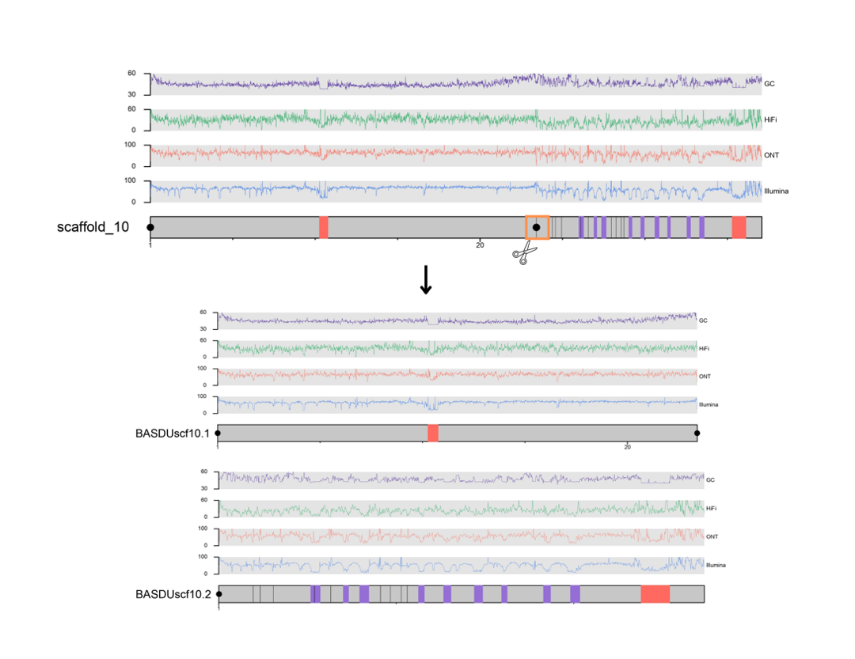

**Figure S2.** Manual breaks arising from internal telomeric sequence. Note that centromeric sequence (red bars, CEN199; purple bars, CEN187) is often associated with a distinct drop in GC content and read depth. Black dots indicate telomeric sequence.

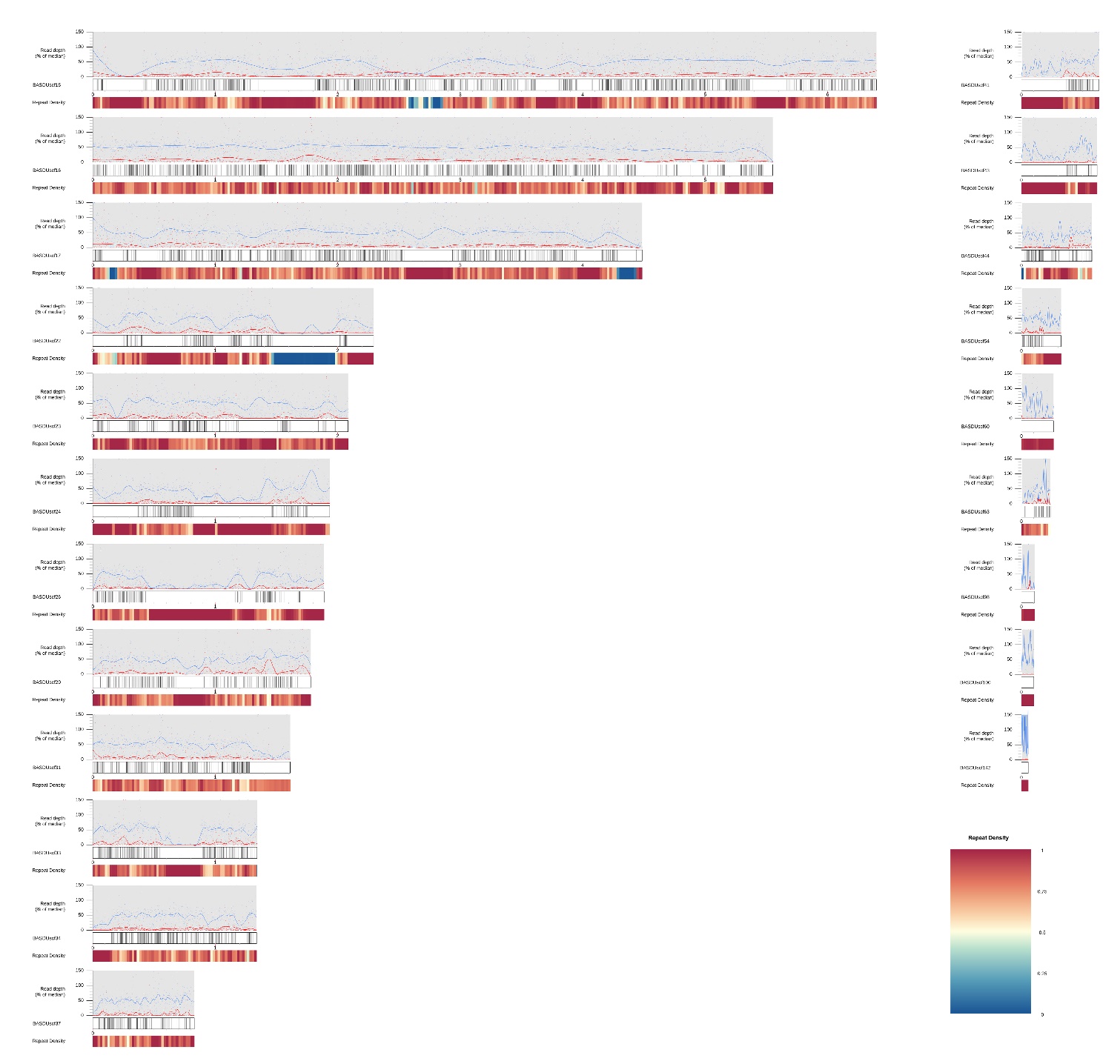

**Figure S3.** Mapping of Y-enriched k-mer contigs to the assembly. Scaffolds with a high density of Y-enriched contigs are putative Y chromosome scaffolds. Shown also is repeat density in 20 kb windows and read depth traces in 1 kb windows. Clearly, the putative Y chromosome is fragmented, with the Y enriched contigs mapping to 21 scaffolds, 11 ≥ 1 Mbp. Refer to the density mapping of Y enriched contigs to autosomes and the X in Figure S4. A high resolution version of this figure can be found at <https://github.com/kango2/basdu>.

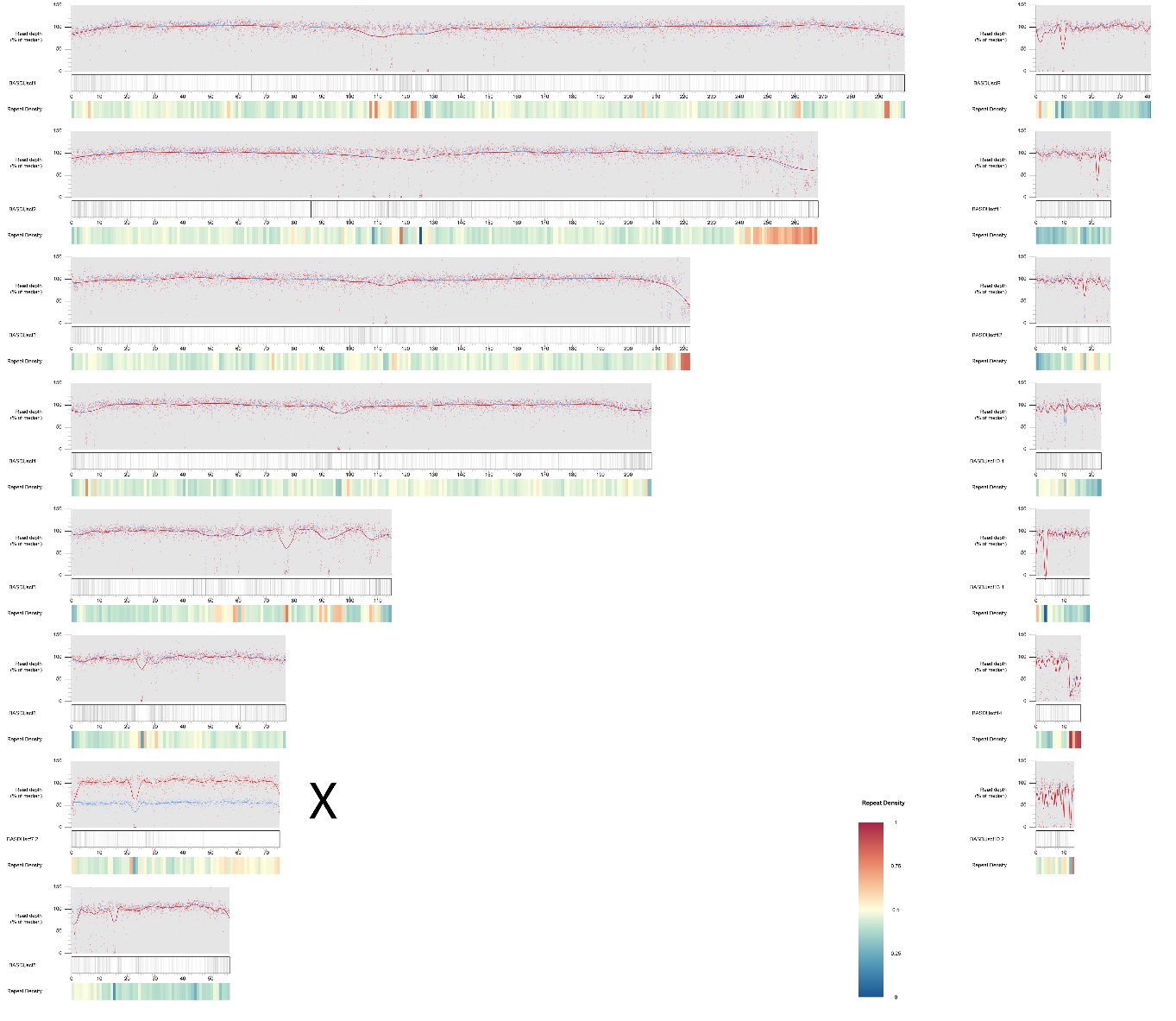

**Figure S4.** A plot of the 15 longest scaffolds (corresponding to the number of chromosomes of *Bassiana duperreyi*) for the *YaHS* assembly. Scaffold BASDUscf7.2 is the X chromosome. Read depth traces in 20kb windows were generated by mapping Illumina genomic reads from a male XY individual (bd65) to the genome assembly (blue), and mapping the Illumina genomic reads from a female XX individual (bd64) to the genome assembly (red). Shown also is repeat density in 1Mb windows. Scaffold BASDUscf7.2 is consistent with an X scaffold that has no homology to the Y chromosome (that is, XX 100% read depth; XY 50% read depth). A high resolution version of this figure can be found at <https://github.com/kango2/basdu>.

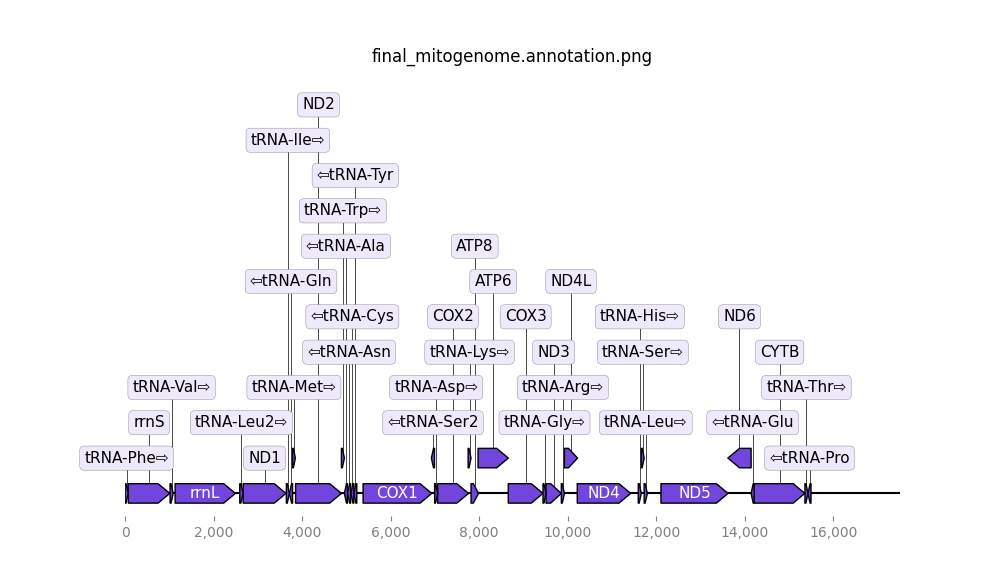

**Figure S5**. Annotation of the mitochondrial genome of *Bassiana duperreyi* using *mitoHiFi*. Control region not shown. Length 17,506 bp.

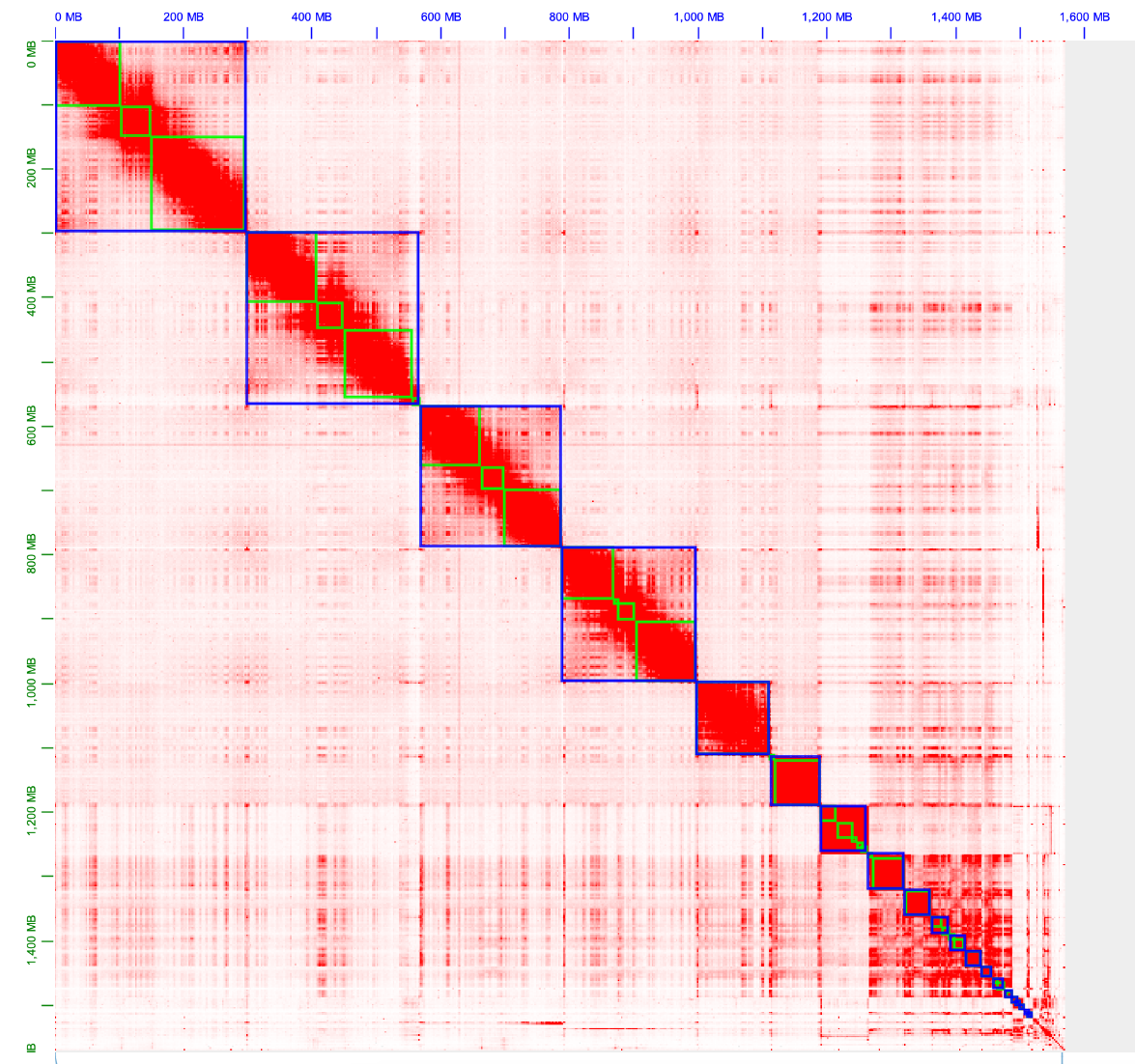

**Figure S6.** HiC contact maps displayed using Juicer for the curated assembly. Ordered by size.
